# Plasticity and lineage commitment of individual Th1 cells are determined by stable T-bet expression quantities

**DOI:** 10.1101/2022.08.14.503916

**Authors:** Ahmed N. Hegazy, Caroline Peine, Dominik Niesen, Isabel Panse, Yevhen Vainshtein, Christoph Kommer, Qin Zhang, Tobias M. Brunner, Michael Peine, Anja Fröhlich, Naveed Ishaque, Roman M. Marek, Jinfang Zhu, Thomas Höfer, Max Löhning

## Abstract

T helper 1 (Th1) cell identity is defined by the expression of the lineage-defining transcription factor T-bet. Here, we examine the influence of T-bet expression heterogeneity on subset plasticity by leveraging cell sorting of distinct *in vivo*-differentiated Th1 cells based on their quantitative expression of T-bet and interferon-γ. Heterogeneous T-bet expression states were regulated by virus-induced type-I interferons and were stably maintained even after secondary viral infection. Exposed to Th2-polarizing conditions, the sorted subpopulations exhibited graded levels of plasticity: T-bet quantities were inversely correlated with the ability to express the Th2 lineage-specifying transcription factor GATA-3 and Th2 cytokines. Reprogramed Th1 cells acquired graded, but stable mixed Th1+2 phenotypes with a hybrid epigenetic landscape. Continuous presence of T-bet in differentiated Th1 cells was essential to ensure Th1 cell stability. Thus, innate cytokine signals regulate Th1 cell plasticity via an individual cell-intrinsic rheostat to enable T cell subset adaptation to subsequent challenges.

**HIGHLIGHTS:**

- Type-I interferons triggered by infection determine T-bet expression states in Th1 cells
- T-bet and IFN-γ expression states indicate the plasticity of individual Th1 cells
- Individual T-bet expression states and plasticity persist after secondary infection
- Reprogramming yields stable Th1+2 phenotypes and a mixed epigenetic landscape

## INTRODUCTION

Cell fate commitment and terminal differentiation are key processes underlying immunological maturation and functional specialization. T cell differentiation is guided by environmental cues, lineage-specifying transcription factors, and epigenetic modifications. The concept of binary cell fate decision promoted the initial paradigm of Th1 and Th2 cell differentiation being mutually exclusive and highly stable (Abbas et al., 1996; Delbrück, 1949; Gardner et al., 2000; MONOD and JACOB, 1961; Mosmann et al., 1986; Zhou et al., 2012). Lineage-specifying transcription factors, such as T-bet in CD4^+^ T-helper-1 (Th1) cells and GATA-3 in Th2 cells, safeguard long-term lineage stability by maintaining their own expression via transcriptional autoactivation (Afkarian 2002,Hofer 2002,Mullen 2001,Ouyang 2000). Indeed, continuous GATA-3 expression in Th2 cells is needed to maintain the Th2 cell phenotype and prevent reprogramming towards Th1 cell fate (Pai et al., 2004; Zhu et al., 2004)

However, cellular differentiation is not necessarily a terminal decision. Differentiated cells retain all genetic information required to regenerate the whole cellular repertoire of an organism (GURDON et al., 1958; Hochedlinger and Jaenisch, 2002). Global analyses of epigenetic marks in fully differentiated T cell subsets revealed that gene loci encoding key transcription factors of other lineages are often retained in a transcriptionally bivalent, non-silenced state (Wei et al., 2009). In line with these observations, we and others recently observed substantial plasticity in CD4 T cell lineages (Bonelli et al., 2014; Hegazy et al., 2010; Zhu et al., 2010). T cell plasticity was shown to provide functional specialization and adaptation advantages for CD4 T cells to allow fine-tuned effective immune responses (Zhou et al., 2009).

In addition to the general cell fate plasticity, T cell lineage differentiation is accompanied by substantial intrapopulation heterogeneity (Marshall et al., 2011; O’garra et al., 2011; Openshaw et al., 1995) This heterogeneity was made visible by improved technologies for single-cell analysis of both RNA and protein expression, and cannot be explained by genetic variability (Papalexi and Satija, 2018). Indeed, recent findings point to the capacity of T cells to acquire individual quantitative cytokine memory, i.e., they commit to distinct production amounts of a given cytokine that are stably maintained throughout the memory phase (Helmstetter et al., 2015) However, how this intrapopulation heterogeneity is established and if it contributes to the decision-making between T cell plasticity versus stable lineage commitment has remained unresolved.

To address the link between intrapopulation heterogeneity and plasticity states, we employed lymphocytic choriomeningitis virus (LCMV) infections to generate LCMV-specific Th1 cells *in vivo* and analyzed their plasticity. Here, we sorted *in vivo*-differentiated Th1 cells by their expression amounts of T-bet and the Th1 signature cytokine interferon-γ (IFN-γ). When stimulated in the presence of Th2-polarizing cytokines, the sorted Th1 cell subpopulations revealed distinct susceptibility to reprogramming: the higher the T-bet expression state, the lower the cell’s plasticity. Hence, Th1 cells with relatively low T-bet expression states featured high expression quantities of GATA-3 and Th2 cytokines after reprogramming. Indeed, the observed intrapopulation heterogeneity and plasticity was a stable feature of the Th1 cell subpopulations generated *in vivo* during viral infections. Presence of T-bet in already committed Th1 cells was required to maintain Th1 cell stability and prevent acquisition of Th2 cell characteristics. The hybrid Th1+2 phenotype of reprogramed Th1 cells was mirrored in corresponding epigenetic landscapes. We thus provide evidence that intrapopulation heterogeneity in expression states of a master-regulator transcription factor determines cell fate decisions of individual cells, thereby explaining plasticity as well as stable lineage commitment within a given cell subset.

## RESULTS

### *In vivo*-primed Th1 cells maintain plasticity towards Th2 program

To generate *in vivo*-primed Th1 cells, a potent Th1 cell-favoring environment was induced in mice by LCMV infection (LCMV, strain WE, 200 PFU; Zinkernagel et al., 1986, Löhning et al., 2008). Naïve TCR-transgenic LCMV-specific CD4^+^ T cells were adoptively transferred into C57BL/6 mice (WT mice), which were then infected with LCMV (Figure 1A). Analysis of the donor cells at the peak of the immune response revealed rigorous expansion of LCMV-specific CD4^+^ T cells with a CD44^hi^CD62L^lo^ effector/memory phenotype (Figure S1A, B). LCMV-induced effector cells expressed T-bet (Figure 1B) as well as the canonical Th1 markers CXCR3 and IL-18Rα and high levels of IFN-γ, TNF-α, and IL-2 (Figure S1C, D). They did not express GATA-3, RORγt, BCL6, FoxP3, or any canonical Th2, Th9, Th17, or Tfh surface markers or cytokines (Figure 1B and S1C, D, E). Similar to T cell receptor (TCR)-transgenic LCMV-specific CD4^+^ T cells, endogenous GP_61-80—_specific effector CD4^+^ T cells, identified by the expression of CD40L upon re-stimulation with dominant GP_61-80_ epitope, exhibited similar T-bet expression and Th1 differentiation (Figure S1F, G).

**Figure 1.**
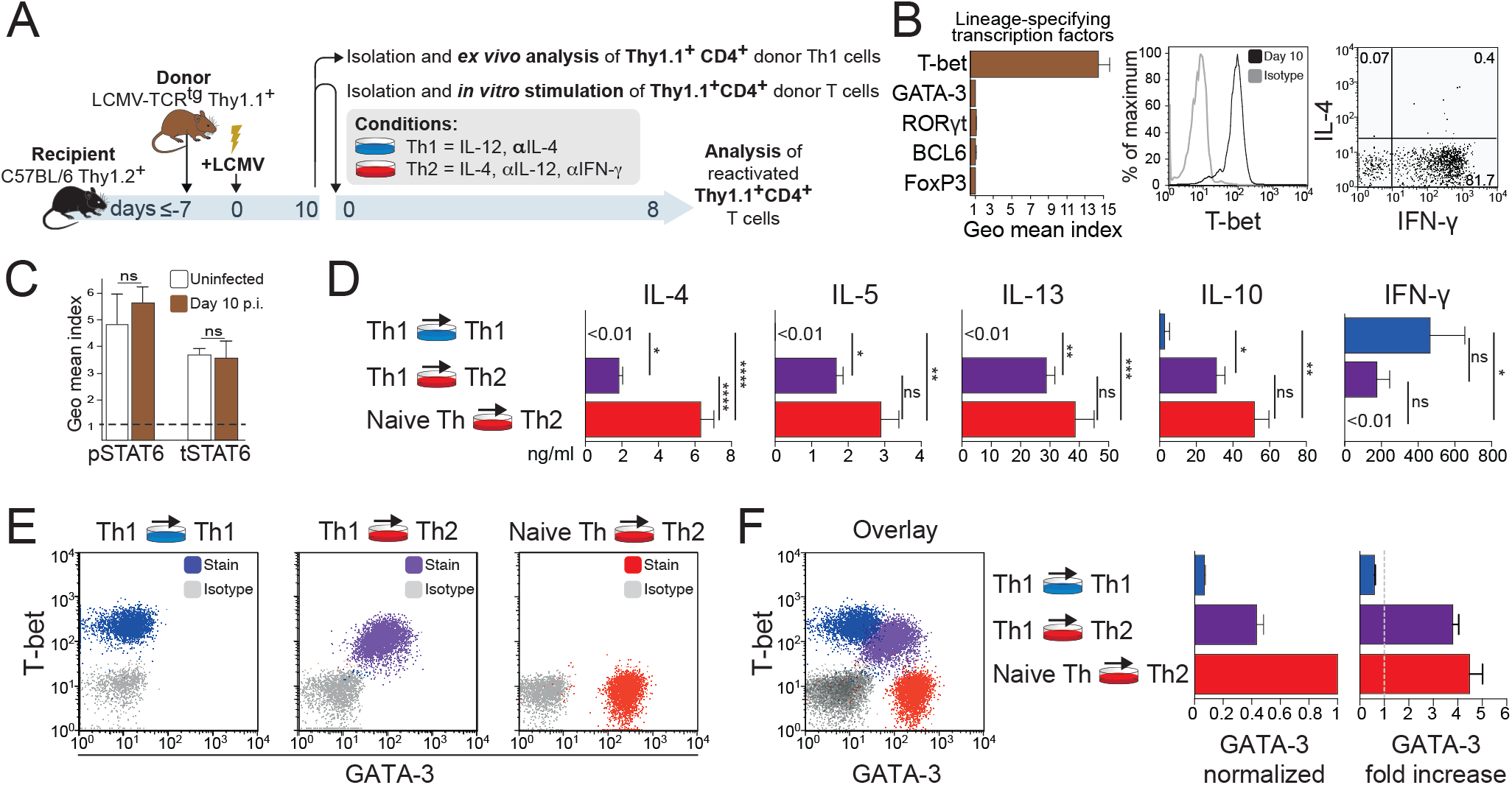
*In vivo*-primed Th1 cells maintain plasticity towards Th2 program. (A-F) Naïve LCMV-specific CD4^+^ Thy1.1^+^ cells were adoptively transferred into normal C57BL/6 mice that were then infected with LCMV. ns, not significant; * *P* < 0.05; ** *P* < 0.01; *** *P* < 0.001; **** *P* < 0.0001. (A) Experimental scheme demonstrating the adoptive cell transfer procedure and analysis time points after infection. (B) 10 days after infection, expression of the indicated lineage-specifying transcription factors and signature cytokines were determined in splenic CD4^+^ Thy1.1^+^ donor cells by FACS. Shown are geometric mean indices +/-SEM of T-bet (Th1), GATA-3 (Th2), RORγt (Th17), BCL6 (Tfh), or FoxP3 (Treg) protein expression in CD4^+^ Thy1.1^+^ donor cells versus endogenous naïve CD62L^hi^ CD44^lo^ CD4^+^ Thy1.2^+^ T cells. Dot plot showing IFN-γ and IL-4 expression after PMA/ionomycin stimulation of splenic CD4^+^ Thy1.1^+^ donor cells. (C) IL-4–induced STAT6 phosphorylation and total STAT6 protein levels in naïve (uninfected) and effector splenic CD4^+^ Thy1.1^+^ donor cells. Geometric mean indices + SEM of pSTAT6 staining of cytokine-exposed vs. untreated cells (left bar graph) and of total STAT6 staining vs. isotype control staining (right bar graph) within CD4^+^ Thy1.1^+^ donor cells are depicted. (D) Cytokine production of CD4^+^ Thy1.1^+^ donor cells treated as indicated was measured upon re-stimulation for 24 h with PMA/ionomycin using cytometric bead arrays. (E, F) GATA-3 and T-bet protein expression of CD4^+^ Thy1.1^+^ donor cells, reactivated as described above, were determined by FACS. (F) Dot plot overlays illustrate GATA-3 and T-bet co-expression. GATA-3 relative protein expression is depicted after normalization to classic Th2 cells (left bar graph). Relative increase in GATA-3 protein expression is plotted compared to levels of GATA-3 in naïve T cells or Th1 cells (right bar graph). (A-F) Data are pooled from three independent experiments (n=9).

Next, we investigated Th1 cell plasticity by sorting *in vivo*-primed Th1 cells from the infected hosts and subjecting them to specific differentiation signals (Figure 1A). When reactivated under classic Th17-, regulatory T cell (Treg)-, or Th9-inducing conditions, Th1 cells failed to express the relevant lineage-specifying molecules RORγt, FoxP3, or IL-9, respectively, and maintained T-bet and IFN-γ expression (Figure S2A-C). However, *in vivo*-primed Th1 cells responded to IL-4 by phosphorylating STAT6 (Figure 1C) and up-regulated several classic Th2-related cytokine genes at both the RNA and protein levels (Figure 1D and S3A). While these Th1 cells up-regulated GATA-3 expression, they still maintained expression of T-bet and IFN-γ (Figure 1E, F). Similar to the TCR-transgenic LCMV-specific CD4^+^ T cells, endogenous Th1-differentiated LCMV-specific CD4^+^ T cells acquired GATA-3 expression upon IL-4 treatment, confirming the above results achieved using TCR-transgenic effector CD4^+^ T cells (Figure S3B).

Interestingly, IL-4 treatment of *in vivo* LCMV-primed Th1 cells achieved intermediate GATA-3 induction compared to Th2 cells polarized from naïve CD4^+^ T cells (Figure 1F, left graph). However, when quantified, the fold increase in GATA-3 expression in IL-4–treated Th1 cells was similar to that observed after induction of classic Th2 cells from naïve CD4^+^ T cells (Figure 1F, right graph) due to lower GATA-3 expression in *in vivo*-primed Th1 cells compared to naïve CD4^+^ T cells.

Thus, *in vivo*-induced Th1 cells selectively up-regulate GATA-3 expression and give rise to a hybrid T-bet^+^GATA-3^+^ Th1+2 phenotype upon IL-4 treatment. However, the molecular signature and chromatin landscape of such a hybrid phenotype were subject of the following experiments.

### *In vivo*-primed Th1 cells acquire an additional Th2-like chromatin landscape upon IL-4 challenge

To gain insight into the epigenetic adaptation during the acquisition of Th2 features in *in vivo*-differentiated Th1 cells, ChIP-seq was performed to globally map histone modifications associated with active (H3K27ac, H3K4me3, and H3K4me1) and repressed (H3K27me3 and H3K9me3) genes. Th1 cells were isolated on day 10 of LCMV infection and subjected to stimulation under either neutral or Th2-favoring (i.e., IL-4) conditions (Figure 2A). Naïve CD4^+^ T cells were compared with *in vivo*-primed Th1 cells, IL-4–treated *in vivo*-primed Th1 cells, and classic *in vitro*-differentiated Th2 cells.

**Figure 2.**
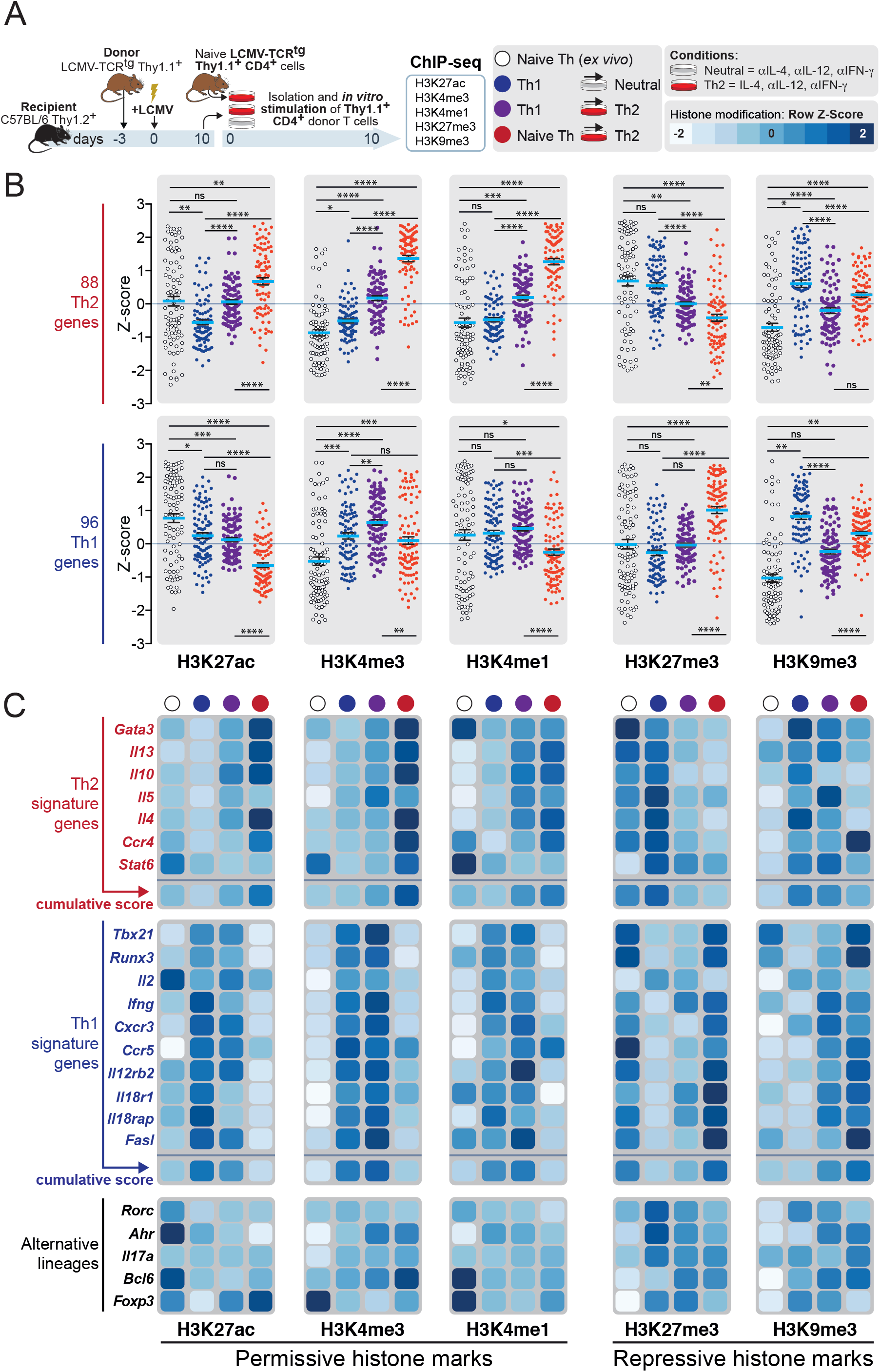
IL-4 drives *in vivo*-primed Th1 cells to acquire an additional Th2-like epigenetic imprint. (A-C) Naïve LCMV-specific CD4^+^ Thy1.1^+^ cells were adoptively transferred into C57BL/6 mice, which were subsequently infected with LCMV. Effector Th1 Thy1.1^+^ donor T cells were isolated on day 10 of infection and reactivated for two rounds of 5 days under neutral (anti–IL-4, anti–IL-12, and anti–IFN-γ) or Th2 (IL-4, anti–IL-12, and anti–IFN-γ) conditions. ns, not significant; * *P* < 0.05; ** *P* < 0.01; *** *P* < 0.001; **** *P* < 0.0001. (A) Experimental scheme. (B, C) H3K27ac, H3K4me3, H3K4me1, H3K27me3, and H3K9me3 modifications of selected Th2 and Th1 genes are presented. (B) Z-scores of all 88 Th2 or 96 Th1 genes of each respective histone modification of the different cell types are depicted. (C) Histone modifications in individual characteristic Th2 or Th1 signature genes in the cell types indicated at the top of the panels are plotted. The color-coding depicts the Z-score value. A Z-score equal to 0 means that the score for a particular gene is the same as the mean for all conditions, while a Z-score of +1 indicates that the corresponding value is one standard deviation above the mean. (B-C) Data represent one experiment with two independent biological replicates pooled from 4-8 independent mice.

To assess the epigenetic status, clusters of genes known to show lineage-specific expression were identified and analyzed (Wei et al., 2009, 2011). *In vivo*-primed Th1 cells and classic *in vitro*-differentiated Th2 cells showed a high density of permissive histone marks (H3K27ac, H3K4me3, and H3K4me1) at Th1- and Th2-related genes, respectively. Consistent with this, repressive marks (H3K27me3 and H3K9me3) densely populated the opposing lineage-related genes (Figure 2B, C). IL-4–treated Th1 cells often showed unaltered permissive marks at Th1-related genes and intermediate permissive and repressive marks at Th2-related genes when compared with classic Th1 and Th2 cells (Figure 2B, C).

These data show that IL-4 treatment superimpose a Th2-like epigenetic imprinting in *in vivo*-differentiated Th1 cells, thus generating a dual Th1 and Th2 chromatin landscape. However, the Th2 chromatin landscape in hybrid T-bet^+^GATA-3^+^ Th1+2 cells exhibited intermediate permissive and repressive marks reflecting a cell-intrinsic regulation of functional plasticity in Th1 cells.

### Type I interferons restrain maximal GATA-3 induction in Th1 cells

We then sought to evaluate the role of critical Th1-inducing signals as well as T-bet in regulating Th1 cell plasticity in viral infections. Therefore, we adoptively transferred naive LCMV-specific CD4^+^ T cells of either fully functional control genotype or with deficiencies in T-bet (*Tbx21*^*-/-*^) or in receptors for IFN-γ (*Ifngr1*^*-/-*^) or type I IFNs (*Ifnar1*^*-/-*^) into wild-type (WT) recipients or IL-12–deficient mice. Then we infected all groups with LCMV, and subsequently isolated LCMV-specific effector CD4^+^ T cells (Figure 3A). All donor-derived effector LCMV-specific CD4^+^ T cells, including donor cells primed in IL-12–deficient mice, with the exception of *Tbx21*^*-/-*^ T cells, acquired a Th1 cell phenotype and normal activation marker expression (Figures 3B-D and S4A, B). However, expression of T-bet and the Th1-associated markers CXCR3 and Ly6C were significantly lower in LCMV-specific CD4^+^ T cells lacking the receptor for type I IFNs (Figure 3B-D). In comparison, type I IFN receptor-competent cell populations expressed approximately twice as much T-bet protein and twice as many cells expressed the Th1-associated markers, indicating that for Th1 lineage differentiation in LCMV infection context, IFN-γ or IL-12 signals are of minor importance. The reduced T-bet expression magnitude in type I IFN receptor-deficient Th1 cells was accompanied by lower amounts of IFN-γ production per cell, whereas the frequency of IFN-γ producing cells was not affected (Figure S4B). *Tbx21*^*-/-*^ CD4^+^ T cells expressed little IFN-γ and failed to express CXCR3 and Ly6C (Figures 3B-D and S4B). Noteworthy, although *Ifnar1*^*-/-*^ Th1 cells featured reduced amounts of T-bet, they did not produce Th2 or Th17 cytokines *ex vivo* (Figure S4C). Thus, type I interferons represent an essential external signal for optimal expression of T-bet and Th1 signature genes during viral infection.

**Figure 3.**
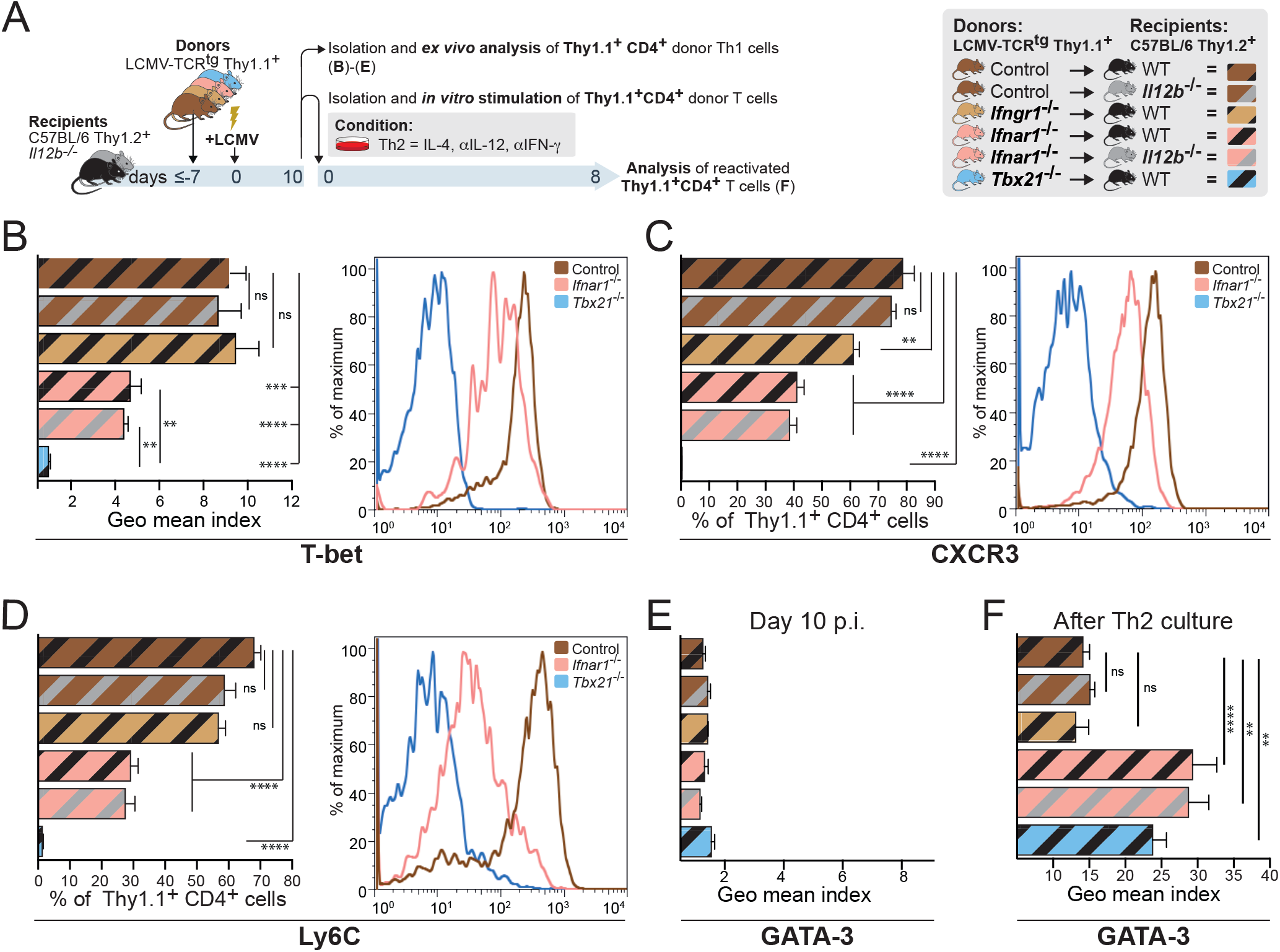
Type I interferons restrain maximal GATA-3 induction in Th1 cells. (A-F) Naïve WT, *Ifnar1*^*-/-*^, *Ifngr1*^*-/-*^, and *Tbx21*^*-/-*^ LCMV-specific CD4^+^ Thy1.1^+^ cells were adoptively transferred into C57BL/6 mice or *Il12b*^*-/-*^ mice. Recipient mice were infected with LCMV. On day 10 after LCMV infection, CD4^+^ Thy1.1^+^ donor T cells were isolated, characterized, and reactivated for two rounds of 4 days under Th2 conditions. ns, not significant; ** *P* < 0.01; *** *P* < 0.001; **** *P* < 0.0001. (A) Experimental scheme. (B) T-bet protein expression was determined in splenic CD4^+^ Thy1.1^+^ donor cells by FACS on day 10 p.i.. Geometric mean indices + SEM of T-bet staining within CD4^+^ Thy1.1^+^ donor T cells are depicted. (C, D) CD4^+^ Thy1.1^+^ donor T cells were analyzed by FACS for expression of the indicated surface molecules on day 10 p.i.. (E, F) Geometric mean indices + SEM of GATA-3 within CD4^+^ Thy1.1^+^ donor T cells isolated directly after infection (day 10 p.i.) or after reactivation for two rounds of 4 days under Th2 conditions. (A-F) Data are pooled from three independent experiments (n=6-13).

Intrigued by the above findings that type I interferons are required for optimal T-bet induction in Th1 cells, we reactivated the same genotype combinations of *in vivo*-generated Th1 cells under Th2-polarizing conditions (Figure 3E, F). Although *ex vivo* all groups featured low GATA-3 levels (Figure 3E), after Th2 culture those cells with deficiencies in receptors for type I IFNs (*Ifnar1*^*-/-*^) or in T-bet (*Tbx21*^*-/-*^) displayed about double the amount of GATA-3 protein than control or *Ifngr1*^*-/-*^ T cells (Figure 3F). However, GATA-3 up-regulation was not observed when the isolated T cells were maintained under neutral conditions (Figure S4D), excluding a default mechanism in *Ifnar1*^*-/-*^ and *Tbx21*^*-/-*^ T cells to up-regulate GATA-3. Thus in *in vivo*-induced *Ifnar1*^*-/-*^ and *Tbx21*^*-/*^ Th1 cells, IL-4 signals up-regulate GATA-3 expression and give rise to a GATA-3^high^ profile in contrast to wild-type Th1 cells.

Taken together, type I interferons maximize T-bet expression during viral infection and stabilize Th1 cell differentiation by restricting GATA-3 induction upon subsequent IL-4 signaling. Hence, we hypothesized that T-bet expression quantity regulates the extent of Th1 cell plasticity.

### T-bet acts in a dose-dependent manner to stabilize the Th1 program and repress the Th2 program

To examine the effect of graded T-bet expression in regulating Th1 cell plasticity, we adoptively transferred naïve *Tbx21*^*+/+*^, *Tbx21*^*+/-*^, and *Tbx21*^*-/-*^ LCMV-specific CD4^+^ T cells into C57BL/6 mice, infected the recipients with LCMV, and analyzed effector CD4^+^ T cells on day 10 post infection (Figure S5A). As previously reported, we observed a dose-dependent reduction in T-bet expression and IFN−γ production in *Tbx21*^*+/-*^ and *Tbx21*^*-/-*^ CD4^+^ T cells compared to *Tbx21*^*+/+*^ cells after infection (Figure 4A and (Marshall et al., 2011; Szabo et al., 2002)). However, the reduction or absence of T-bet did not influence GATA-3 expression after LCMV infection (Figure 4A).

**Figure 4.**
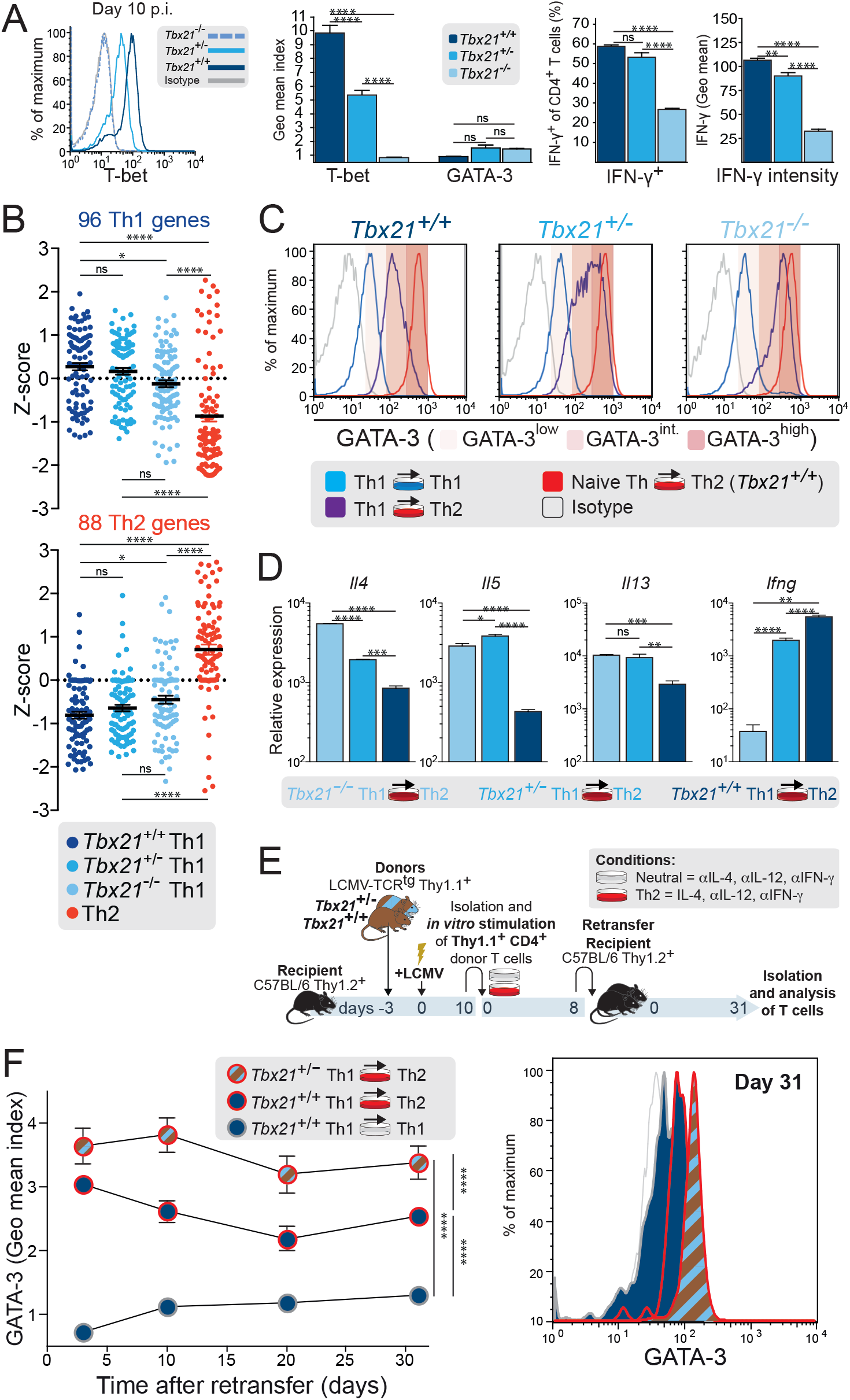
T-bet acts in a dose-dependent manner to stabilize the Th1 program and repress the Th2 program. (A-D) Naïve *Tbx21*^*+/+*^, *Tbx21*^*+/-*^, and *Tbx21*^*-/-*^ LCMV-specific CD4^+^ Thy1.1^+^ cells were adoptively transferred into C57BL/6 mice. Recipient mice were infected with LCMV. ns, not significant; * *P* < 0.05; ** *P* < 0.01; *** *P* < 0.001; **** *P* < 0.0001. (A) After 10 days of infection, T-bet and GATA-3 protein expression were determined in splenic CD4^+^ Thy1.1^+^ donor cells by FACS. Geometric mean indices + SEM of T-bet and GATA-3 staining within CD4^+^ Thy1.1^+^ donor T cells are depicted (histogram and middle bar graphs). IFN-γ production of CD4^+^ Thy1.1^+^ donor cells was measured upon re-stimulation of spleen cells with PMA/ionomycin (right bar graphs). Mean frequencies and geometric mean + SEM of IFN-γ^+^ within CD4^+^ Thy1.1^+^ donor cells are depicted. (B) RNA-seq analysis of selected 96 Th1 and 88 Th2 genes in *in vivo*-primed *Tbx21*^*+/+*^, *Tbx21*^*+/-*^, and *Tbx21*^*-/-*^ LCMV-specific CD4^+^ Thy1.1^+^ cells after expansion *in vitro* under neutral conditions. Data represent one experiment with two independent biological replicates pooled from 7-9 independent mice. (C, D) CD4^+^ Thy1.1^+^ donor T cells of the indicated genotypes were isolated and reactivated for two rounds of 5 days under Th1 (IL-12, anti–IL-4, and anti–IFN-γ) or Th2 conditions. Th2 cells derived from WT naïve LCMV-specific CD4^+^ Thy1.1^+^ cells served as control. (C) GATA-3 protein amounts were determined in CD4^+^ Thy1.1^+^ donor cells by FACS. (D) Quantitative PCR of the indicated genes after re-stimulation with PMA/ionomycin for 3 h. Data are pooled from three independent experiments (n=6-9). (E-F) Naïve *Tbx21*^*+/+*^ and *Tbx21*^*+/-*^ LCMV-specific CD4^+^ Thy1.1^+^ cells were adoptively transferred into C57BL/6 mice. Recipient mice were infected with LCMV. 10 days after infection, CD4^+^ Thy1.1^+^ donor T cells were isolated and reactivated for two rounds of 4 days under neutral or Th2 conditions. Subsequently, the cells were retransferred into naïve C57BL/6 mice. (E) Experimental scheme. (F) At the indicated time points after retransfer, GATA-3 protein amounts were determined in the peripheral blood of CD4^+^ Thy1.1^+^ cell recipients by FACS. The geometric mean index depicts the factor of change of the GATA-3 geometric mean in CD4^+^ Thy1.1^+^ donor cells compared with the GATA-3 values in endogenous naive CD62L^hi^CD4^+^Thy1.1^-^ T cells. Data represent the mean ± SEM of n = 4-6 mice per group and time point. ****p < 0.001 at all measured time points.

To evaluate the effect of T-bet dose on the transcriptional and epigenetic landscape of Th1 cells, we performed transcriptomic and epigenetic analysis by RNA-seq and modification-specific histone ChIP-seq on *in vivo*-primed *Tbx21*^*+/+*^, *Tbx21*^*+/-*^, and *Tbx21*^*-/-*^ LCMV-specific CD4^+^ T cells. To assess the transcriptional and epigenetic status, Th1- and Th2-specific clusters of genes described in Figure 2 were analyzed (Wei et al., 2009, 2011). T-bet dose did not influence either the levels of permissive marks (H3K27ac, H3K4me3, H3K4me1) or repressive marks (H3K27me3, H3K9me3) at Th1-related genes in *Tbx21*^*+/-*^ versus *Tbx21*^*+/+*^ CD4^+^ T cells (Figure S5B). However, complete T-bet loss in *Tbx21*^*-/-*^ CD4^+^ T cells led to a substantial reduction in permissive marks and an increase in repressive marks at Th1-related genes, even though the cells were primed in the same Th1-favoring conditions as the other genotypes. These epigenetic patterns were reflected by the expression behavior of the Th1-related gene cluster (Figure 4B). As previously reported, *Tbx21*^*-/-*^ CD4^+^ T cells were biased toward Th2 and Th17 phenotypes (Intlekofer et al., 2008; Zhu et al., 2012) represented by an increased expression of Th2 signature genes (Figure 4B) as well as a decrease in repressive marks and an increase in permissive marks at Th2-related genes and the Th17-related genes *Rorc* and *Il17a* when compared with T-bet–competent CD4^+^ T cells (Figure S5B).

Of note, reduced T-bet expression in *Tbx21*^*+/-*^ CD4 T cells had a stepwise influence on repressive histone modifications at Th2-related gene loci, especially on H3K27me3, without significant influence on gene expression of Th2-related genes (Figures S5B and 4B). Thus, maximal T-bet expression is not only essential for maximal accessibility of Th1 genes but is also important for repression of Th2- and Th17-associated genes such as *Gata3, Il13, Il5, Il4, Rorc*, and *Il17a* in Th1 cells. Indeed, *Tbx21*^*+/-*^ CD4^+^ T cells expressed higher levels of GATA-3 and Th2 cytokines compared to *Tbx21*^*+/+*^ CD4^+^ T cells after culturing under Th2 conditions (Figure 4C, D). In line with this, IL-4 treatment induced the highest GATA-3 levels in *Tbx21*^*-/-*^ CD4^+^ T cells (Figure 4C).

Taken together, these data highlight the role of T-bet dose in the epigenetic imprinting of the Th1 lineage and its stability against alternative differentiation signals.

### Long-term maintenance of graded GATA-3 expression in Th1 cells

Based on our observations, Th1 cells can acquire at least three levels of GATA-3 expression: classic Th1 cells are low or negative for GATA-3 but, after IL-4 treatment, intermediate GATA-3 levels are induced in *Tbx21*^*+/+*^ Th1 cells and higher levels in *Tbx21*^*+/-*^ Th1 cells. We next sought to assess the stability of graded GATA-3 levels in IL-4–treated Th1 cells.

To achieve graded induction of GATA-3 expression in Th1 cells, *Tbx21*^+/+^ and *Tbx21*^+/-^ LCMV-specific naive CD4^+^ T cells were adoptively transferred into WT mice and primed by LCMV infection. These *in vivo*-generated Th1 cells were then isolated, subjected to Th2-differentiating conditions or kept under neutral conditions, and adoptively retransferred into uninfected WT mice (Figure 4E). We tracked GATA-3 and T-bet protein expression of the transferred cells for more than a month. Compared with Th1 cells kept under neutral conditions, IL-4-treated *Tbx21*^+/-^ Th1 cells showed the highest expression quantity of GATA-3, while IL-4-treated *Tbx21*^+/+^ Th1 cells reached an intermediate amount. This relative GATA-3 distribution was maintained throughout the entire time-course and was reflected in the homogeneous peaks of the cell populations from the distinct genotypes (Figure 4F). In both genotypes, T-bet expression quantities were kept at equal levels between IL-4-primed and classic Th1 cells, indicating that the reprogramming did not diminish T-bet expression intensity (Figure S5C, right). Graded GATA-3 levels correlated with distinct expression capacities for Th2 but not Th1 cytokines (Figure S5D). The graded GATA-3 expression was maintained not only at steady state in resting memory T cells but also after T cell reactivation with cognate antigen *in vitro* (Figure S5E). These results demonstrate the long-term stability of T-bet–regulated graded GATA-3 expression states in hybrid T-bet^+^ GATA-3^+^ Th1+2 cells.

### The magnitude of T-bet and IFN-γ expression predicts Th1 cell plasticity

Previously we demonstrated intrapopulation heterogeneity of Th1 cells, with single Th1 memory cells recalling the production of individual IFN-γ and T-bet amounts (Helmstetter et al., 2015). To address the impact of this intrapopulation heterogeneity on Th1 cell plasticity, we utilized transgenic mice expressing a fluorescent reporter protein, ZsGreen, under the control of the T-bet promoter(Zhu et al., 2012). These *T-bet-ZsGreen* (TBGR) mice were infected with LCMV, and activated CD4^+^CD44^hi^CD62L^lo^ T cells were sorted 10 days after infection according to ZsGreen (ZsG) expression intensities (Figure 5A). ZsG expression quantitatively correlated with T-bet and IFN-γ protein production (Figure 5A, B). GATA-3 levels were low in all sorted ZsG populations (Figure 5B). When reactivated in Th2-polarizing conditions, all sorted fractions up-regulated GATA-3 protein levels to some extent. However, T-bet-ZsG^lo^- and T-bet-ZsG^int^-sorted cells displayed higher GATA-3 levels than T-bet-ZsG^hi^-sorted Th1 cells, and a substantial fraction of the former T-bet-ZsG^lo/int^ cells even reached the GATA-3^high^ level found in classic Th2 cells (Figure 5C).

**Figure 5.**
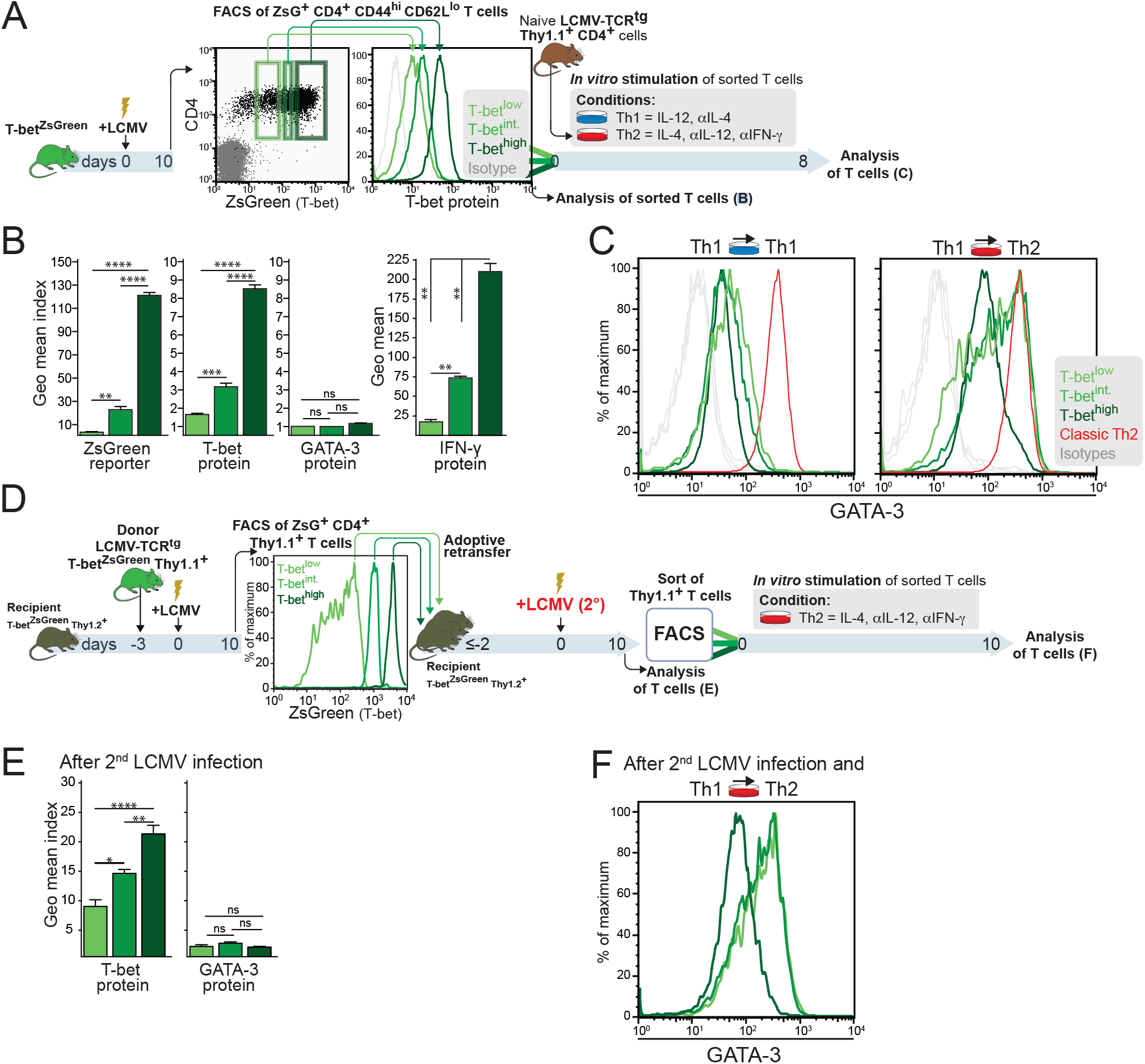
Graded T-bet expression is a stable property of Th1 cells and predicts Th1 cell plasticity. (A-C) Naive *T-bet-ZsGreen* (TBGR) mice were infected with LCMV. 10 days after infection, CD4^+^ CD44^hi^ CD62L^lo^ T cells were sorted according to ZsGreen expression intensity by FACS. (A) Experimental scheme. (B) ZsGreen intensity and T-bet and GATA-3 protein expression were determined in sorted T cell fractions by FACS. (B, left bar graphs) Geometric mean indices + SEM of ZsGreen intensity and T-bet and GATA-3 staining of the sorted T cell populations are indicated. Geometric mean + SEM of IFN-γ expression intensity within gated ZsG^hi^, ZsG^int^, and ZsG^lo^ CD4^+^ Thy1.1^+^ donor cells was analyzed after re-stimulation with GP_61-80_ by intracellular cytokine staining. ns, not significant; * *P* < 0.05; ** *P* < 0.01; *** *P* < 0.001; **** *P* < 0.0001. (C) Endogenous LCMV-specific CD4^+^ T cells were reactivated under the indicated conditions with GP_61-80_ and APCs. Th2 cells derived from naïve LCMV-specific CD4^+^ cells served as control. GATA-3 protein expression was determined in activated CD4^+^ CD154^+^ T cells by FACS. (A, C) Data are pooled from two independent experiments with three independent biological replicated derived from n=20. (D-E) Naïve LCMV-specific TBGR CD4^+^ Thy1.1^+^ cells were adoptively transferred into untreated TBGR Thy1.2^+^ mice. Recipient mice were infected with LCMV. Effector CD4^+^ Thy1.1^+^ T cells were FACS sorted according to ZsGreen expression intensity. Equal numbers of isolated CD4^+^ ZsG^+^ Thy1.1^+^ cells were retransferred into uninfected TBGR Thy1.2^+^ recipient mice that were subsequently infected with LCMV. ns, not significant; * *P* < 0.05; ** *P* < 0.01; *** *P* < 0.001; **** *P* < 0.0001. Data are pooled from two independent experiments with three independent biological replicated derived from n=20. (D) Experimental scheme. (E) Ten days after secondary infection, T-bet and GATA-3 protein expression of donor CD4^+^ Thy1.1^+^ T cells was determined. (F) CD4^+^ Thy1.1^+^ donor T cells were isolated after secondary infection and reactivated *in vitro* under Th2 conditions. GATA-3 protein expression was analyzed by FACS.

Since the usage of a T-bet reporter is restricted to transgenic cells, we next employed naturally occurring markers that correlate with T-bet expression quantities. Previously we found that individual Th1 cells feature stable graded expression amounts of T-bet that correlate with the quantity of secreted IFN-γ (Helmstetter et al., 2015). We therefore sort-purified LCMV-specific Th1 cells, generated *in vivo* as shown in Figure 1A, with distinct IFN-γ secretion intensities (IFN-γ^hi^, IFN-γ^int^, and IFN-γ^lo^ fractions, Figure S6A) and tested their plasticity under Th2-polarizing conditions. Here, the majority of the IFN-γ^lo^ Th1 cells reached GATA-3 amounts of classic Th2 cells, whereas this was achieved by approximately half of the IFN-γ^int^ cells and only few cells of the IFN-γ^hi^ fraction (Figure S6B). Enhanced GATA-3 induction was accompanied by elevated Th2 cytokine production (Figure S6C).

Taken together, purification of Th1 cells according to T-bet expression intensities or IFN-γ production amounts separates flexible (T-bet^int/lo^) from stably committed (T-bet^hi^) Th1 cells. These results suggest that T-bet quantity regulates the degree of plasticity of *in vivo*-primed Th1 cells. Thus, the generation of a gradient of T-bet expression during primary anti-viral CD4^+^ T cell responses maintains functional plasticity as well as stability in this differentiated T cell population.

### Graded T-bet expression is a stable property of Th1 cells

To study whether the above-observed gradient of T-bet expression persists after secondary infection, we purified T-bet^hi^, T-bet^int^, and T-bet^lo^ Thy1.1^+^ LCMV-specific CD4^+^ T cells according to ZsG expression intensities after primary infection with LCMV. Equal numbers of graded ZsG-sorted populations were then retransferred into uninfected recipient mice (Figure 5D), and ZsG and intracellular T-bet protein expression were quantified after secondary LCMV infection (Figure 5E).

Graded T-bet expression levels established during primary infection were maintained after secondary infection (Figure 5E), arguing against incomplete differentiation of the subpopulations with intermediate or low T-bet expression during primary infection. Instead, intrapopulation heterogeneity of T-bet quantities appears to be an inherent feature of virus-specific Th1 cells.

Next, we compared the ability of T-bet-ZsG^hi^, T-bet-ZsG^int^, and T-bet-ZsG^lo^ Thy1.1^+^ LCMV-specific CD4^+^ T cells to respond to IL-4 treatment after secondary infection. Sorted T-bet^lo^ and T-bet^int^ T cells acquired higher levels of GATA-3 expression compared to T-bet^hi^ T cells, similar to the functional plasticity seen in these Th1 cell subpopulations after primary infection (Figure 5F, compare with Figure 5C).

In conclusion, differential T-bet expression states are stably imprinted in individual Th1 cells during primary CD4^+^ T cell responses. These graded T-bet expression quantities generate functional heterogeneity within the antigen-specific CD4^+^ T cell pool to allow subsequent adaption of Th1 responses to changes of the immunological microenvironment.

### Continuous T-bet expression safe-guards Th1 cell stability

We demonstrated above that T-bet acts in a dose-dependent manner to stabilize the Th1 differentiation program while repressing Th2 differentiation. However, in the above settings, T-bet levels were either genetically altered from the beginning or were imprinted to be expressed at low levels during the initial priming. To evaluate whether the presence of T-bet in already committed Th1 cells is required to maintain Th1 cell stability, we used a conditional deletion approach where LCMV-specific cells were differentiated normally into Th1 cells *in vivo* during LCMV infection and only then we induced the deletion of one or both *Tbx21* alleles by tamoxifen treatment. To this end, we adoptively transferred naive *Tbx21*^*fl/fl*^ and *Tbx21*^*fl/+*^ Rosa26CreERT2^+^ LCMV-specific CD4^+^ T cells into WT mice, which were subsequently infected with LCMV (Figure 6A). Mice were then treated with tamoxifen, and afterwards we analyzed the LCMV-specific Thy1.1^+^ CD4^+^ T cells (Figure 6A). All donor-derived T cell populations exhibited an activated Th1 cell phenotype (Figures 6B and S7A). However, T-bet protein amounts were slightly reduced in *Tbx21*^fl/+^ T cells and strongly diminished in *Tbx21*^*fl/fl*^ T cells (Figure 6C). The reduction or absence of T-bet did not influence GATA-3 expression in these cells after LCMV infection (Figure 6C).

**Figure 6.**
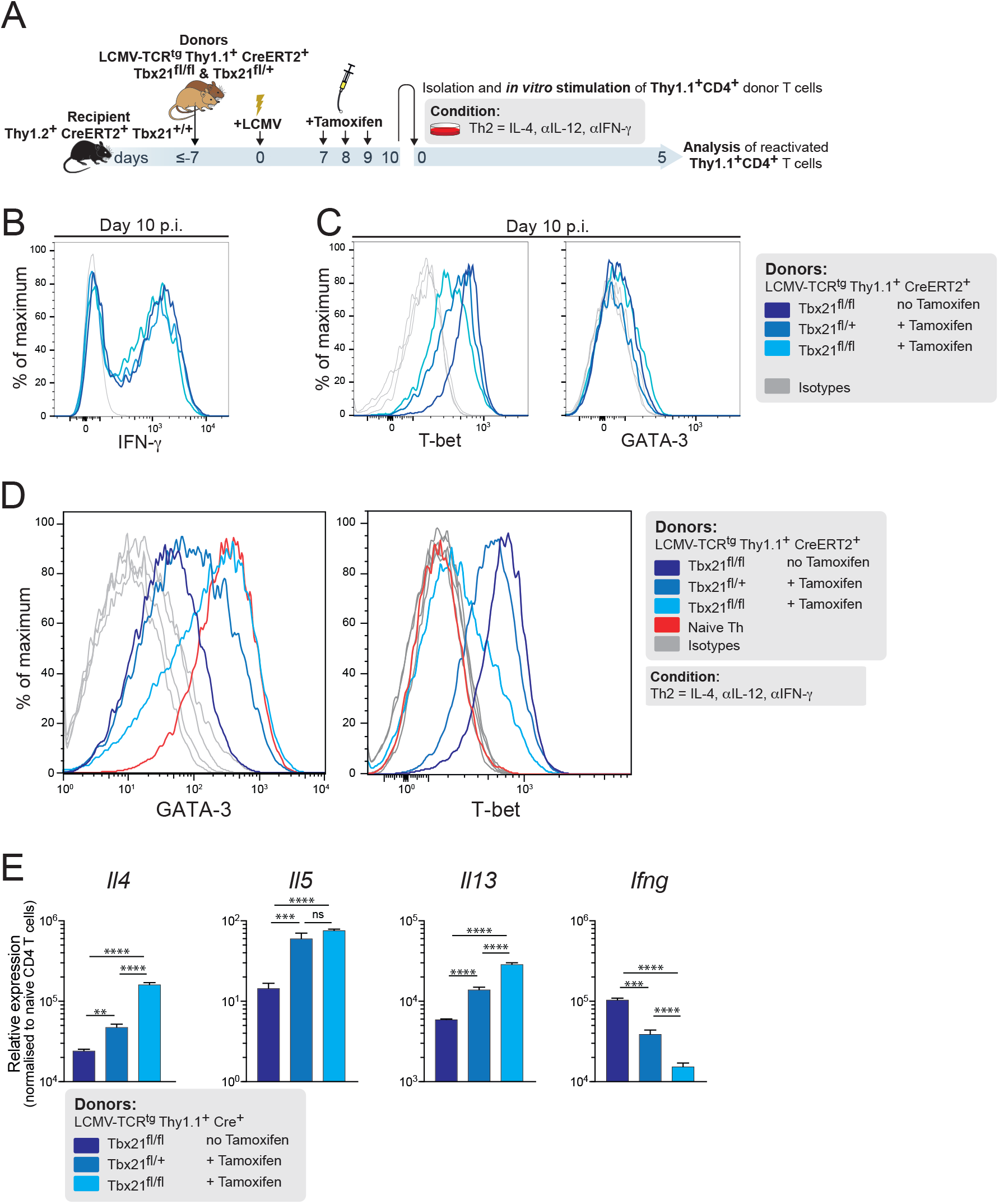
Continuous T-bet expression safe-guards Th1 cell stability. Naïve *Tbx21*^*fl/fl*^ and *Tbx21*^*fl/+*^ Rosa26CreER^T2+^ LCMV-specific CD4^+^ Thy1.1^+^ cells were adoptively transferred into Th1.2^+^ Rosa26CreER^T2+^ C57BL/6 mice. Recipient mice were infected with LCMV. On day 7, 8, and 9 p.i., recipient mice were treated with tamoxifen while a control group was left untreated. On day 10 p.i., CD4^+^ Thy1.1^+^ donor T cells were isolated and directly analyzed or reactivated for 5 days under Th2 conditions. Th2 cells derived from naïve LCMV-specific CD4^+^ Thy1.1^+^ cells served as control. ns, not significant; ** *P* < 0.01; *** *P* < 0.001; **** *P* < 0.0001. Data are representative of two independent experiments. (A) Experimental scheme. (B) IFN-γ expression and (C) T-bet and GATA-3 expression were determined in CD4^+^ Thy1.1^+^ donor cells by FACS on day 10 p.i.. (D) GATA-3 and T-bet protein expression was determined by FACS in CD4^+^ Thy1.1^+^ donor cells after re-activation for 5 days under Th2 conditions. (E) Quantitative PCR of the indicated cytokine genes after re-stimulation with PMA/ionomycin for 3 h.

We then sorted the LCMV-primed Thy1.1^+^ Th1 cells of the above genotypes from the tamoxifen-treated or untreated recipients and tested their plasticity. When reactivated in Th2-polarizing conditions, but not in Th1 or neutral conditions (Figure S7B), the sorted tamoxifen-treated genotypes up-regulated GATA-3 protein levels to a certain degree when compared to the *Tbx21*^*fl/fl*^ T cells without tamoxifen treatment (Figure 6D). However, tamoxifen-treated *Tbx21*^*fl/fl*^ T cells reached GATA-3 amounts of classic Th2 cells, whereas *Tbx21*^*fl/fl*^ T cells from tamoxifen-untreated recipients (T-bet-sufficient T cells) showed intermediate GATA-3 protein amounts. Furthermore, the enhanced GATA-3 induction in Th1 cells after deletion of one or both T-bet alleles was accompanied by gradually elevated Th2 cytokine production and graded reduction in IFN-γ expression (Figure 6E). These results demonstrate that the graded presence of T-bet in differentiated Th1 cells is required to regulate the extent of Th2 cell reprogramming competence.

## DISCUSSION

Here we have addressed Th1 cell lineage commitment and plasticity in light of intrapopulation heterogeneity. Our findings provide insight into the molecular network controlling cell fate decisions in individual Th1 cells. We have identified specific environmental signals and lineage-specifying transcription factors in conjunction with epigenetic changes as driving forces of cellular differentiation and functionality. Importantly, we found stable individual quantitative expression states of the lineage-specifying transcription factor T-bet at the core of Th1 intrapopulation heterogeneity. In Th1 cells generated *in vivo* during viral infection, type I IFNs are required to achieve maximal T-bet expression. While being fully differentiated, T-bet^lo/int^ Th1 cells are more susceptible to reprogramming: When subjected to Th2 signals, they start to express higher levels of GATA-3 and Th2 cytokines and acquire additional Th2-like epigenetic features. The lower the T-bet expression state, the higher is the cell’s plasticity. Thus, intrapopulation heterogeneity in expression states of a master regulator transcription factor explains why stable lineage commitment and plasticity can be observed in a Th1 cell population at the same time (Figure S8). The consideration of quantitative aspects reconciles seemingly contradictory findings of previous studies on Th1 cell lineage commitment (Abbas et al., 1996; Assenmacher et al., 1998; DuPage and Bluestone, 2016; Krawczyk et al., 2007; Mosmann et al., 1986; Murphy and Stockinger, 2010; Murphy et al., 1996; Nakayamada et al., 2012; Panzer et al., 2011; Perez et al., 1995).

The importance of quantitative aspects in the expression of key transcription factors and cytokines has emerged only relatively recently (Helmstetter et al., 2015; Marshall et al., 2011; O’garra et al., 2011) We here show that type I IFNs imprint an individual T-bet expression state in a developing Th1 cell during its primary activation *in vivo*. This specific T-bet expression state is preserved even when the cell is reactivated by a secondary viral infection, and it is not lost when the cell receives Th2-polarizing signals. Quantitative differences between individual Th1 cells are cell-intrinsically maintained and highly stable both in the effector and memory phase.

How do individual Th1 cells memorize their specific T-bet expression state? We hypothesize that the combination of transcriptional autoactivation of key transcription factors together with modifications of the epigenetic landscape are main players in the continuous process of maintaining a cell’s identity – not only at the qualitative, but also at the quantitative level (Zhou and Huang, 2011). In addition, individual T-bet levels might also influence the cytokine sensitivity of a Th1 cell by regulating its cytokine receptor expression or the downstream transmission of cytokine signals. However, we compared cells with lower T-bet expression states, e.g., Th1 cells with only one functional *Tbx21* allele, with *Tbx21*^*+/+*^ Th1 cells, and found a highly similar expression pattern of both Th1- and Th2-associated genes (compare Figure 4B). Hence, T-bet^lo/int^ cells cannot be regarded as weaker or insufficiently differentiated Th1 cells. Yet, these cells exhibit a higher level of plasticity: They express more GATA-3 and Th2 cytokines when reactivated in a Th2-polarizing environment. Similar quantitative mechanisms of cell fate control might also take place in other cell populations with intrapopulation heterogeneity, e.g., tumor cells (Lu et al., 2013; Ross et al., 2003), embryonic stem cells (Kolodziejczyk et al., 2015), and thrombocytes (Macaulay et al., 2016)

Beyond its classic role as transcription factor directly regulating the transcription of target genes, T-bet enforces lineage commitment in two additional ways. Firstly, T-bet shapes the epigenetic landscape of T cells to induce the expression of Th1-specific genes, which occurs in concert with cytokine signal-transducing STAT proteins (Vahedi et al., 2012). Here, T-bet physically interacts with several chromatin remodeling complexes and recruits them to its target genes (Kanno et al., 2012; Wei et al., 2009; Zhu and Paul, 2010) Secondly, T-bet blocks the expression of genes associated with alternative lineages (Oestreich and Weinmann, 2012). For example, it indirectly represses Th2 or Th17 signature genes by protein-protein interactions with GATA-3 or Runx1, respectively, thereby preventing them from binding to their DNA target sites (Hwang et al., 2005; Lazarevic et al., 2010). We found that reprogramming of Th1 cells to the Th1+2 phenotype resulted in significantly modified histone marks at Th2 gene loci whereas Th1 loci remained mostly unaffected. In a further attempt to explore how T-bet expression influences Th1 stability and plasticity, Kommer et al. (co-submitted manuscript) combined the ChipSeq and RNAseq data generated from T-bet^+/+^, Tbet^+/-^ and Tbet^-/-^ Th1 and Th1+2 cells together with machine learning to study the influence of T-bet expression levels on the epigenetic regulation of Th1 cells. Kommer et al. identified several regulatory enhancer types that fell into three categories: regulated by T-bet, regulated by Th2 cytokines, and controlled by both types of factors (“mixed” enhancers). When analysing genome-wide binding of key Th cell transcription factors, T-bet-regulated enhancers bound T-bet as well as STAT1 and STAT4, while Th2-cytokine-regulated enhancers predominantly bound STAT6 and GATA3. T-bet-regulated enhancer activity was gradually lost with declining Tbx21 gene dose. These findings support our observation of enhanced Th1 cell plasticity in T-bet^int/lo^ Th1 cells and substantiate the relevance of T-bet expression magnitude in maintaining Th1 stability. Thus, our findings pave the way for a better understanding of the regulation of T cell differentiation and plasticity.

The stability of phenotypically distinct populations expanded *in vitro* is of utmost importance for adoptive T cell therapies. In Th1 cells, high T-bet levels are a prerequisite for phenotypic stability. In a non-transgenic setting, intracellular T-bet amounts cannot be detected in living cells. Therefore, we used secreted IFN-γ amounts to identify a Th1 cell’s T-bet expression state. Indeed, the sorted IFN-γ^hi^ cells proved to be most resistant to reprogramming signals. In mice, the intensity of Ly6C expression on the cell surface indicates intracellular T-bet expression amounts (Marshall et al., 2011) and thus it might serve as a surrogate marker for the stability of a Th1 cell.

Our findings have considerable clinical implications. They allow the targeted purification of Th1 cells of defined phenotypic plasticity and the stable functional modulation of these cells for adoptive cell therapies. More generally, we here suggest a model of cell fate decisions in individual cells determined by stable quantitative differences in the expression states of key transcription factors. These differences are caused by differential provision of external cytokine signals. The resulting intrapopulation heterogeneity translates into graded states of lineage commitment and provides the basis for stability or flexibility of individual cells within a population. Reprogramming yields graded hybrid phenotypes with mixed functionality: While the original differentiation program is maintained, additional functions are acquired and imprinted in the epigenetic landscape. Our model of quantitative cell fate determination likely applies to various cell types and may be harnessed for targeted cellular therapies tailored to specific clinical needs.

## EXPERIMENTAL PROCEDURES

### Mice

C57BL/6J were bred under specific pathogen-free conditions at the Charité, Berlin and were used as recipients for adoptive cell transfers at 8-12 weeks. All knockout and SMARTA1 TCR-transgenic (tg) mice, which express an LCMV epitope GP_61–80_-specific TCR, were on the C57BL/6J background (for details, see Supplemental Experimental Procedures). All animal experiments were performed in accordance with the German law for animal protection with permission from the local veterinary offices.

### Adoptive transfer and virus infection

Naive LCMV-specific CD4^+^ T cells were injected intravenously in 500 μl BSS. Mice were infected intravenously with 200 PFU LCMV in 200 μl. For details, see Supplemental Experimental Procedures.

### T cell activation and differentiation

Naïve or effector LCMV-TCR^tg^ CD4^+^ T cells from spleens and lymph nodes were cultured as described previously (Löhning et al., 2008). For details, see Supplemental Experimental Procedures.

### Flow cytometry and cell sorting

Lymphocytes were isolated from peripheral blood, spleens, and lymph nodes and were stained as described previously (Hegazy et al., 2010). Antibody clones and intracellular detection of cytokines, key transcription factors, and STAT proteins are described in the Supplemental Experimental Procedures.

### ChIP-seq and quantitative PCR

ChIP-seq experiments were performed as described previously (Barski et al., 2007; Wei et al., 2009). A detailed description of Chip-seq, RNA-seq and data analysis is provided in the Supplemental Experimental Procedures. For quantitative PCR, RNA was purified using the RNAeasy Mini Kit (Qiagen) and reverse transcribed (Applied Biosystems). Taqman Master Mix and primers from Applied Biosystems were used.

### Statistical analysis

Two groups were compared using the two-tailed unpaired Student’s *t*-test. More than two groups were compared using one-way ANOVA with Bonferroni’s post-hoc test for multiple comparisons. Time courses of multiple groups were compared with two-way ANOVA.

## AUTHOR CONTRIBUTIONS

A.N.H. designed experiments, performed *in vitro* and *in vivo* studies, analyzed data, and wrote the manuscript. C.H., D.N., I.P., Q.Z., M.P., T.M.B. and A.F. performed *in vitro* and/or *in vivo* experiments. Y.V, C.K., N.I., and T.H. performed and supervised bioinformatical analyses. R.M.M., J.Z., and W.E.P. provided reagents and advice. M.L. and T.H. supervised the study, designed experiments, and wrote the manuscript.

## ACKNOWLEDGMENTS

We thank Diana Boesel, Vivien Holecska, Tuula Geske, Heidi Hecker-Kia, and Heidi Schliemann for expert technical assistance and the FCCF at DRFZ for cell sorting. This work was supported by the German Federal Ministry of Education and Research (FORSYS and T-Sys), the German Research Foundation (DFG via SFB618, TPC3; SFB650, TP28; LO1542/3-1, LO1542/4-1, LO1542/5-1, and HO02050/4-1), the Volkswagen Foundation (Lichtenberg fellowship to M.L.), and the Willy Robert Pitzer Foundation (Osteoarthritis Research program to M.L.). A.N.H. is supported by a Lichtenberg fellowship and „Corona Crisis and Beyond“ grant by Volkswagen Foundation, a Berlin Institute of Health Clinician Scientist grant, and the DFG (DFG-TRR241-A05, INST 335/597-1). J.Z. is supported by the Division of Intramural Research, NIAID, NIH, USA. T.H., Y.V., and Q.Z. are supported in part by the US National Science Foundation (NSF PHY11-25915).

The authors declare that no competing interests exist.

## Notes

### Competing Interest Statement

The authors have declared no competing interest.

### Summary of Updates

Updated version of the Figures

