## Supplemental figures and methods for "Plasticity and lineage commitment of individual Th1 cells are determined by stable T-bet expression quantities"

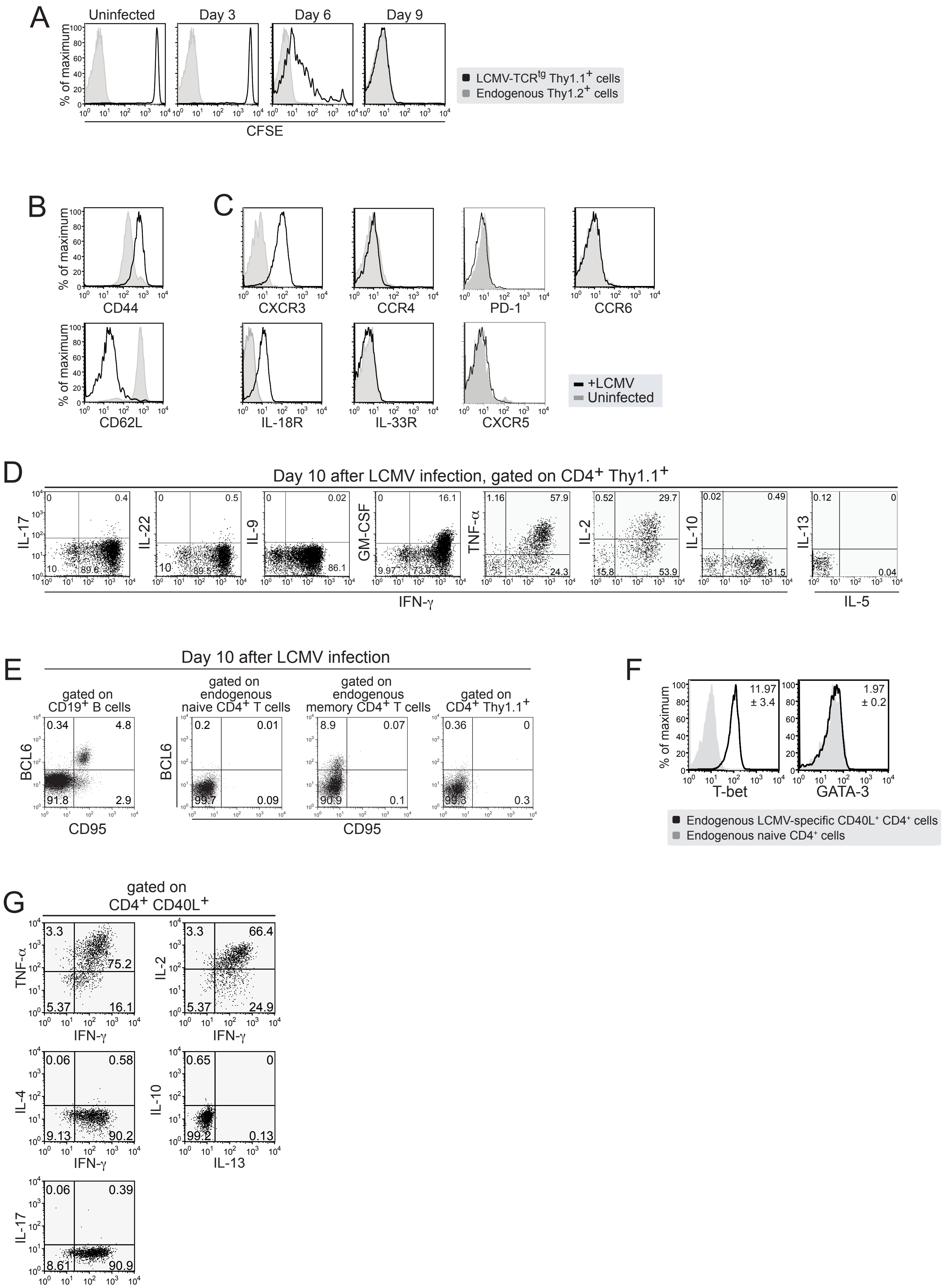
A
Uninfected
Day 3
Day 6
Day 9
% of maximum
CFSE
LCMV-TCR$^{tg}$ Thy1.1$^{+}$ cells
Endogenous Thy1.2$^{+}$ cells
B
C
CD44
CD62L
CXCR3
CCR4
PD-1
CCR6
IL-18R
IL-33R
CXCR5
+LCMV
Uninfected
D
Day 10 after LCMV infection, gated on CD4$^{+}$ Thy1.1$^{+}$
IL-17
IL-22
IL-9
GM-CSF
TNF-α
IL-2
IL-10
IL-13
IFN-γ
IL-5
E
Day 10 after LCMV infection
gated on CD19$^{+}$ B cells
gated on endogenous naive CD4$^{+}$ T cells
gated on endogenous memory CD4$^{+}$ T cells
gated on CD4$^{+}$ Thy1.1$^{+}$
BCL6
CD95
F
T-bet
GATA-3
11.97 ± 3.4
1.97 ± 0.2
Endogenous LCMV-specific CD40L$^{+}$ CD4$^{+}$ cells
Endogenous naive CD4$^{+}$ cells
G
gated on CD4$^{+}$ CD40L$^{+}$
TNF-α
IL-2
IL-4
IL-10
IL-17
IFN-γ
IL-13

Figure S1

**Figure S1. LCMV infection drives strong Th1 cell differentiation and stable T-bet induction**

(A-D) Naïve LCMV-specific $CD4^+$ $Thy1.1^+$ cells were adoptively transferred into C57BL/6 mice, followed by infection of the recipient mice with LCMV. In some experiments, naïve LCMV-specific $CD4^+$ $Thy1.1^+$ cells were labelled with CFSE before adoptive transfer.

(A) CFSE dilution profile of $CD4^+$ $Thy1.1^+$ donor cells was determined at the indicated time points. Data representative of one experiment (n=3-4)

(B, C) Surface expression of the indicated proteins was determined on $CD4^+$ $Thy1.1^+$ donor cells on day 10 after infection.

(D) Ten days after LCMV infection, splenocytes were re-stimulated with PMA/ionomycin, and the frequencies of $cytokine^+$ cells within gated $CD4^+$ $Thy1.1^+$ donor cells were analyzed by intracellular cytokine staining. (C, D) Data representative of three experiments (n=6-9).

(E) Ten days after LCMV infection, splenocytes were stained with BCL6 and CD95 to confirm BCL6 expression in follicular B cells based on our transcription factors staining protocol. Data representative of three independent mice.

(F, G) Naive C57BL/6 mice were infected with LCMV. On day 10 after infection, splenocytes were re-stimulated with $GP_{64-80}$ to reactivate endogenous LCMV-specific $CD4^+$ T cells. Endogenous LCMV-specific $CD4^+$ T cells were identified by CD40L expression. (E) T-bet and GATA-3 expression in $CD4^+$ $CD40L^+$ T cells after $GP_{61-80}$ restimulation were analyzed (black line, stain; gray, isotype; upper panel). (F) Frequencies of $cytokine^+$ cells within gated LCMV-specific $CD4^+$ $CD40L^+$ cells were analyzed by intracellular cytokine staining (lower panel). Data representative of 3-6 experiments.

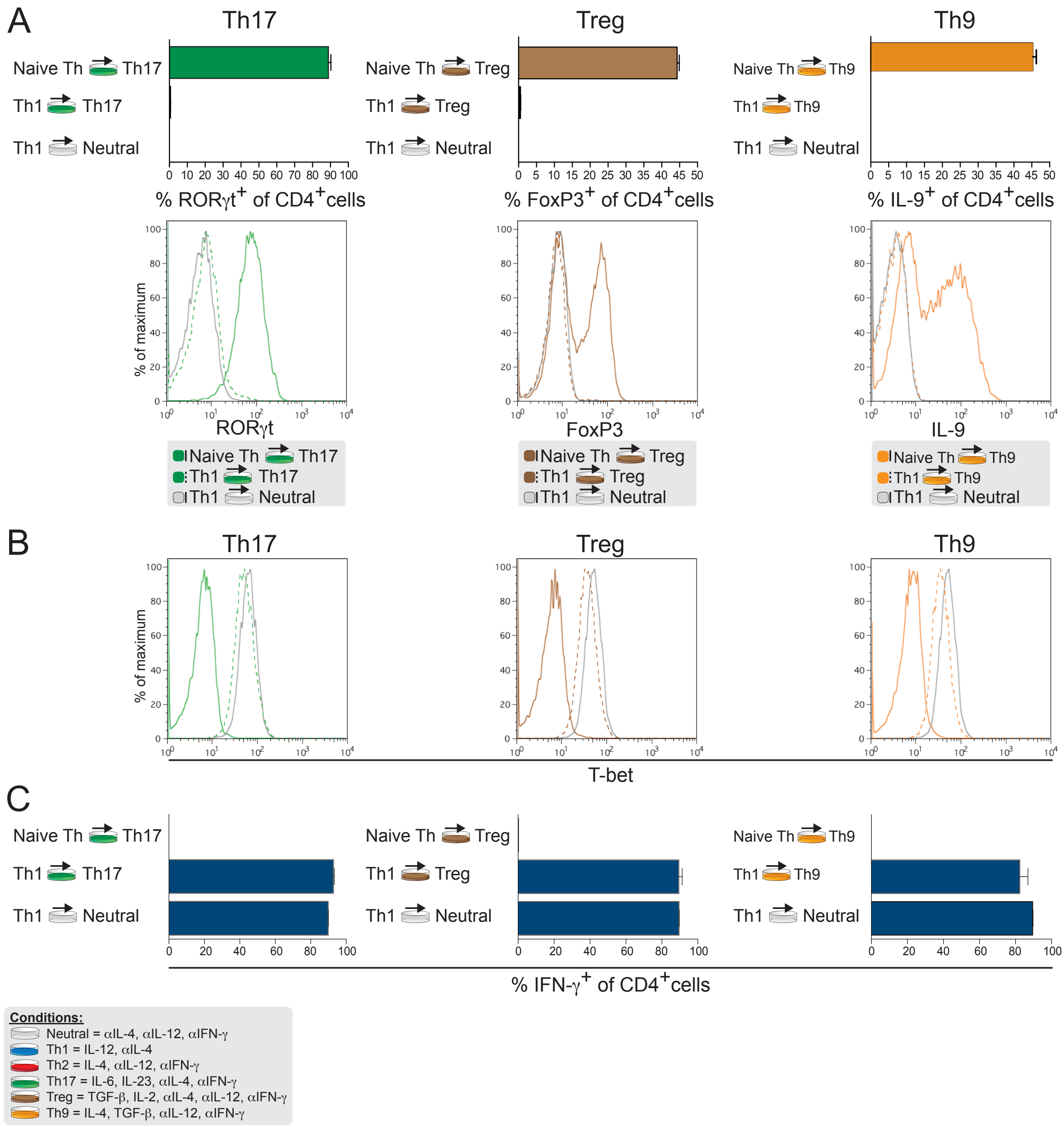
A
Th17
Treg
Th9
Naive Th Th17
Th1 Th17
Th1 Neutral
Naive Th Treg
Th1 Treg
Naive Th Th9
Th1 Th9
% RORγt+ of CD4+cells
% FoxP3+ of CD4+cells
% IL-9+ of CD4+cells
% of maximum
RORγt
FoxP3
IL-9
B
T-bet
C
% IFN-γ+ of CD4+cells
Conditions:
Neutral = αIL-4, αIL-12, αIFN-γ
Th1 = IL-12, αIL-4
Th2 = IL-4, αIL-12, αIFN-γ
Th17 = IL-6, IL-23, αIL-4, αIFN-γ
Treg = TGF-β, IL-2, αIL-4, αIL-12, αIFN-γ
Th9 = IL-4, TGF-β, αIL-12, αIFN-γ

Figure S2

**Figure S2. Plasticity of *in vivo*-primed Th1 cells towards Th17, Treg and Th9 lineages**

(A-C) Naïve LCMV-specific $CD4^+$ $Thy1.1^+$ cells were adoptively transferred into C57BL/6 mice, followed by infection of the recipient mice with LCMV. Effector $CD4^+$ $Thy1.1^+$ donor cells isolated on day 10 after infection were stimulated for 5 days under Th17, Treg or Th9 conditions. WT naïve $CD4^+$ $Thy1.1^+$ cells primed under the same conditions served as a control.

(A) RORγt and FoxP3 expression were assessed on day 5 of culture. For detection of IL-9 production, $CD4^+$ $Thy1.1^+$ cells were re-stimulated on day 5 of cultures with PMA/ionomycin for 4h and then stained for intracellular IL-9.

(B) T-bet expression in $CD4^+$ $Thy1.1^+$ cells was assessed on day 5 of the same cultures described above.

(C) The above-mentioned T cell cultures were re-stimulated with PMA/ionomycin, and the frequencies of IFN-$\gamma^+$ cells within gated $CD4^+$ $Thy1.1^+$ were analyzed by intracellular cytokine staining. Data representative of one experiment (n=3).

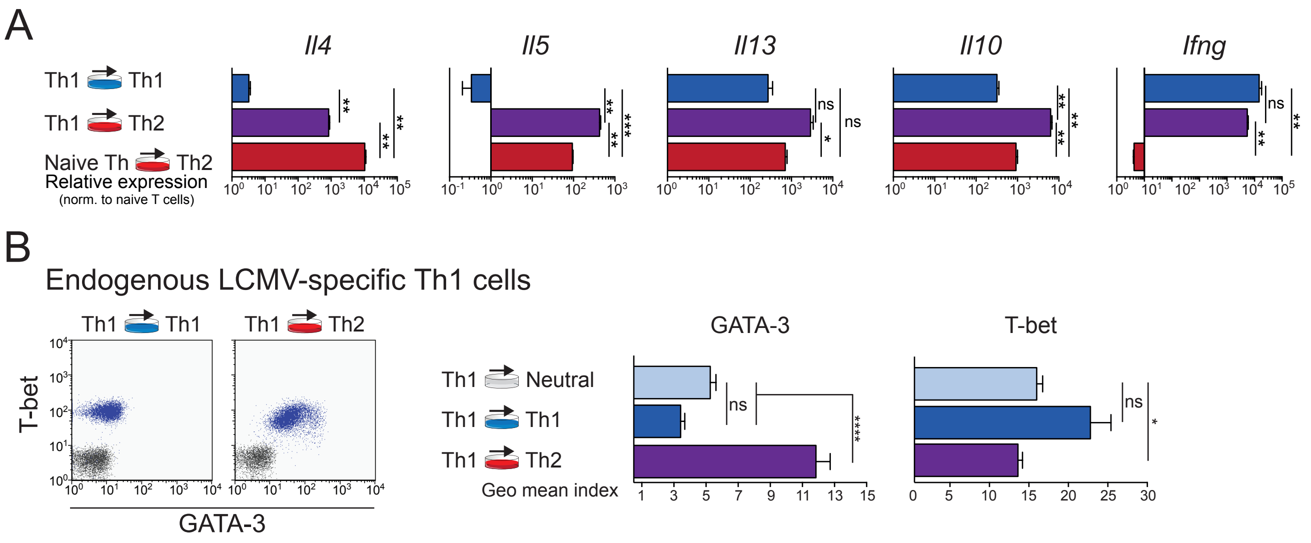
A
Il4
Il5
Il13
Il10
Ifng
Th1 Th1
Th1 Th2
Naive Th Th2
Relative expression
(norm. to naive T cells)
ns
B
Endogenous LCMV-specific Th1 cells
Th1 Th1
Th1 Th2
T-bet
GATA-3
GATA-3
T-bet
Th1 Neutral
Th1 Th1
Th1 Th2
Geo mean index
ns
****
*

Figure S3

**Figure S3. IL-4 induce GATA-3 upregulation in virus-specific Th1 cells**

(A) Naïve LCMV-specific $CD4^+$ $Thy1.1^+$ cells and effector $CD4^+$ $Thy1.1^+$ donor cells isolated on day 10 after infection were reactivated for 4 days under the indicated conditions. Cytokine production of $CD4^+$ $Thy1.1^+$ donor cells was measured upon re-stimulation for 24 hours with PMA/ionomycin using qPCR. Data are representative of three independent experiments (n=6-9).

(B) Naive C57BL/6 mice were infected with LCMV. On day 10 after infection, $CD4^+$ T cells were isolated and cultured under the indicated conditions with $GP_{61-80}$ and APCs to reactivate endogenous LCMV-specific $CD4^+$ T cells. After 4 days, GATA-3 and T-bet protein expression of $CD4^+$ T cells were determined by FACS. Dot plot overlays illustrate GATA-3 and T-bet co-expression. Geometric mean indices ± SEM of GATA-3 or T-bet staining within $CD4^+$ T cells are depicted (bar graphs). Data representative of two experiments.

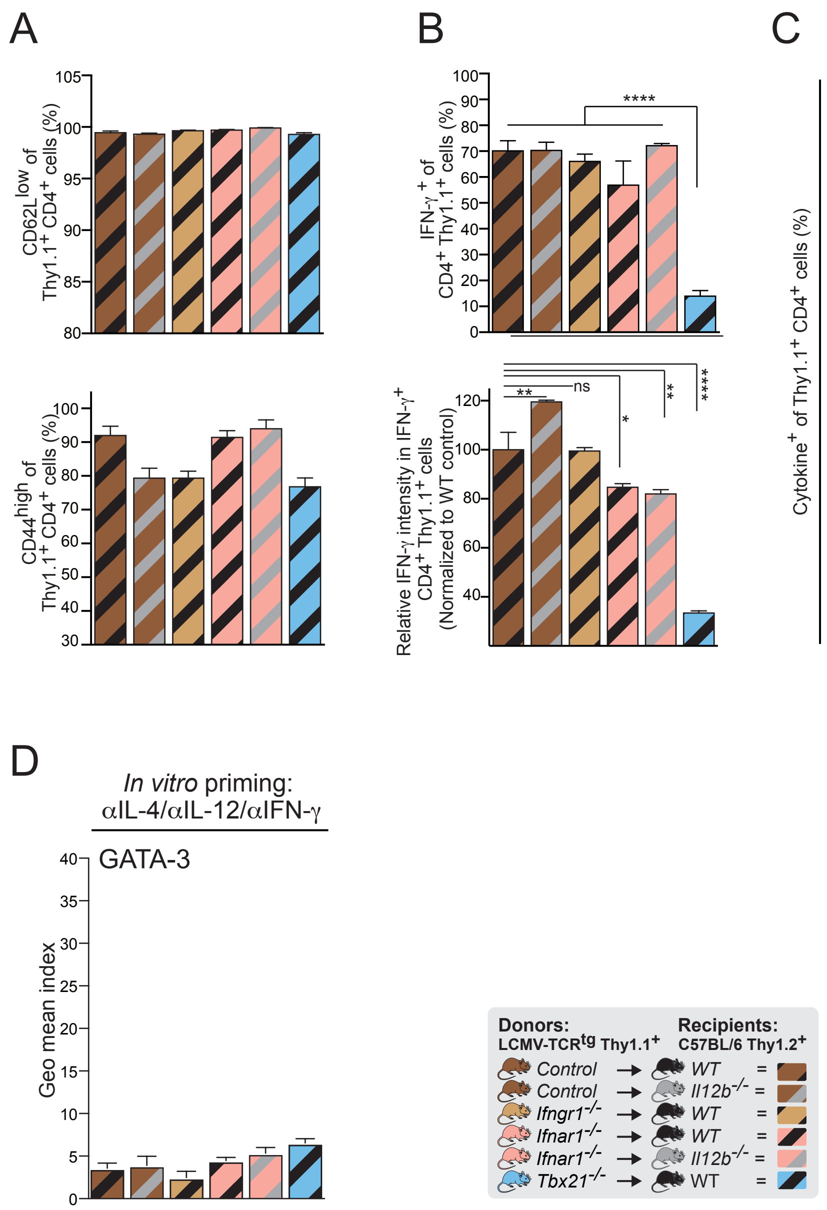
A
CD62L low of Thy1.1+ CD4+ cells (%)
CD44 high of Thy1.1+ CD4+ cells (%)
B
IFN-γ + of CD4+ Thy1.1+ cells (%)
Relative IFN-γ intensity in IFN-γ+ CD4+ Thy1.1+ cells (Normalized to WT control)
ns
C
Cytokine+ of Thy1.1+ CD4+ cells (%)
IL-4
IL-5
IL-13
IL-17
D
In vitro priming:
αIL-4/αIL-12/αIFN-γ
GATA-3
Geo mean index
Donors:
LCMV-TCRtg Thy1.1+
Recipients:
C57BL/6 Thy1.2+
Control → WT =
Control → Il12b-/- =
Ifngr1-/- → WT =
Ifnar1-/- → WT =
Ifnar1-/- → Il12b-/- =
Tbx21-/- → WT =

Figure S4

**Figure S4. Type I interferons regulate maximal T-bet expression and restrain GATA-3 induction in Th1 cells**

(A-C) Naïve WT, *Ifnar1*$^{-/-}$, *Ifngr1*$^{-/-}$, and *Tbx21*$^{-/-}$ LCMV-specific $CD4^+$ Thy1.1$^+$ cells were adoptively transferred into C57BL/6 mice or into *Il12b*$^{-/-}$ mice. Recipient mice were infected with LCMV. Mean frequencies ± SEM within $CD4^+$Thy1.1$^+$ donor cells are indicated. ns, not significant; * $P < 0.05$; ** $P < 0.01$; *** $P < 0.001$.

(A) $CD4^+$ Thy1.1$^+$ T cells were analyzed 10 days after infection by flow cytometry for expression of the indicated surface molecules.

(B, C) On day 10 after LCMV infection, splenocytes were re-stimulated with PMA/ionomycin, and the frequencies of cytokine$^+$ cells within gated $CD4^+$ Thy1.1$^+$ donor cells were analyzed by intracellular cytokine staining.

(D) Geometric mean indices + SEM of GATA-3 within $CD4^+$ Thy1.1$^+$ donor T cells after reactivation for two rounds of 4 days under neutral conditions. (A-D) Data are pooled from three independent experiments (n=6-13).

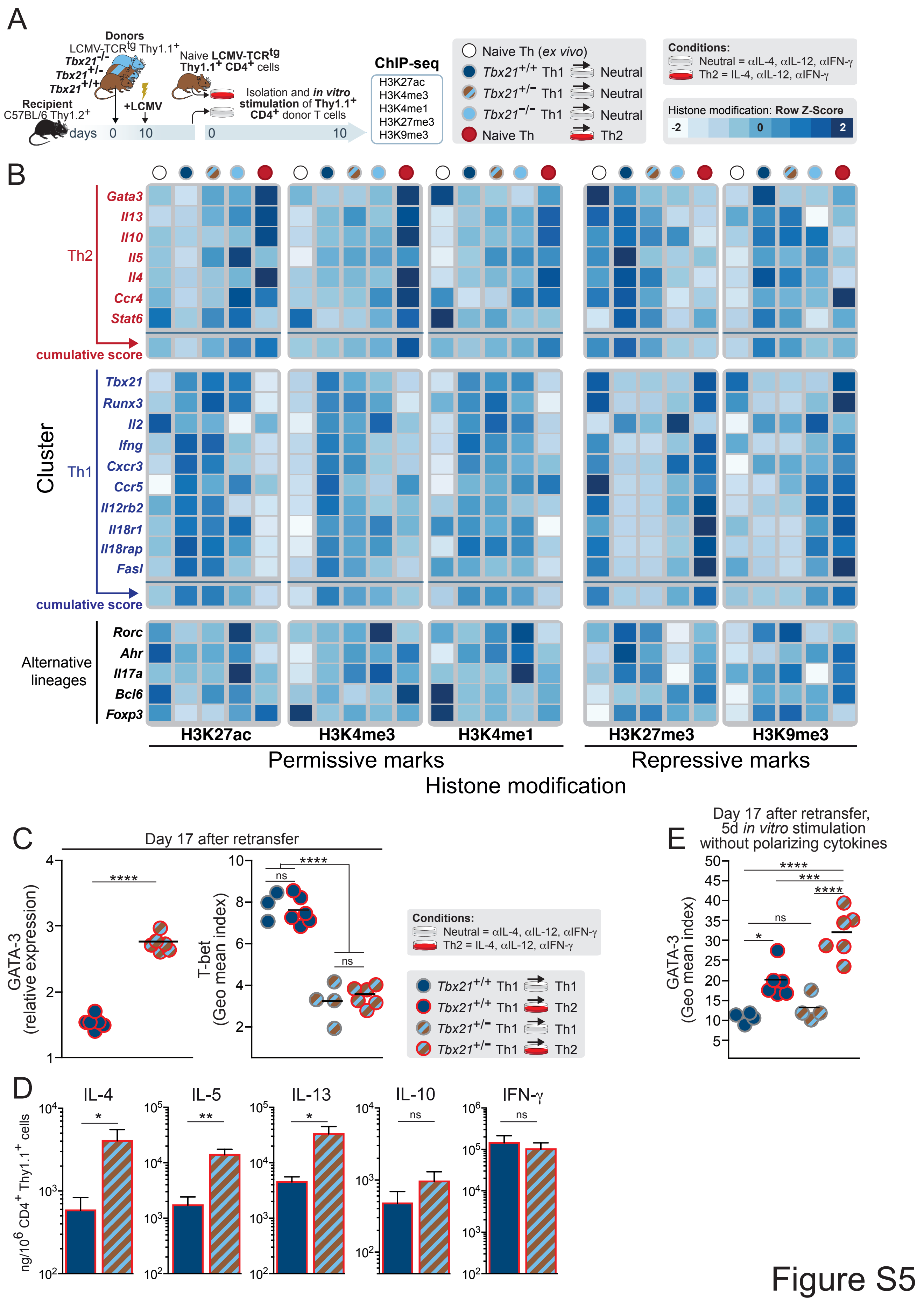

A
Donors
LCMV-TCRtg Thy1.1+
Tbx21-/-
Tbx21+/-
Tbx21+/+
Recipient
C57BL/6 Thy1.2+
+LCMV
days 0 10
Naive LCMV-TCRtg Thy1.1+ CD4+ cells
Isolation and in vitro stimulation of Thy1.1+ CD4+ donor T cells
0 10
ChIP-seq
H3K27ac
H3K4me3
H3K4me1
H3K27me3
H3K9me3
Naive Th (ex vivo)
Tbx21+/+ Th1 Neutral
Tbx21+/- Th1 Neutral
Tbx21-/- Th1 Neutral
Naive Th Th2
Conditions:
Neutral = αIL-4, αIL-12, αIFN-γ
Th2 = IL-4, αIL-12, αIFN-γ
Histone modification: Row Z-Score
-2 0 2
B
Th2
Gata3
Il13
Il10
Il5
Il4
Ccr4
Stat6
cumulative score
Cluster
Th1
Tbx21
Runx3
Il2
Ifng
Cxcr3
Ccr5
Il12rb2
Il18r1
Il18rap
Fasl
cumulative score
Alternative lineages
Rorc
Ahr
Il17a
Bcl6
Foxp3
H3K27ac
H3K4me3
H3K4me1
H3K27me3
H3K9me3
Permissive marks
Repressive marks
Histone modification
C
Day 17 after retransfer
GATA-3 (relative expression)
****
T-bet (Geo mean index)
****
ns
ns
Conditions:
Neutral = αIL-4, αIL-12, αIFN-γ
Th2 = IL-4, αIL-12, αIFN-γ
Tbx21+/+ Th1 Th1
Tbx21+/+ Th1 Th2
Tbx21+/- Th1 Th1
Tbx21+/- Th1 Th2
E
Day 17 after retransfer, 5d in vitro stimulation without polarizing cytokines
GATA-3 (Geo mean index)
****
***
****
ns
*
D
ng/10^6 CD4+ Thy1.1+ cells
IL-4
*
IL-5
**
IL-13
*
IL-10
ns
IFN-γ
ns


Figure S5

**Figure S5. T-bet expression magnitude represses Th2 cytokine genes and permits enhanced GATA-3 induction**

(A, B) Naïve WT, *Tbx21*$^{+/-}$ and *Tbx21*$^{-/-}$ LCMV-specific CD4$^+$ Thy1.1$^+$ cells were adoptively transferred into C57BL/6 mice. Recipient mice were infected with LCMV. ns, not significant; * $P < 0.05$; ** $P < 0.01$; *** $P < 0.001$.

(A) Experimental setup.

(B) Chip-seq analysis of the indicated histone modifications. *In vivo*-primed WT, *Tbx21*$^{+/-}$, and *Tbx21*$^{-/-}$ LCMV-specific CD4$^+$ Thy1.1$^+$ cells were expanded *in vitro* under neutral conditions and then Chip-seq was performed for the indicated histone modifications. The color-coding depicts the Z-score value for each gene based on a scale from -2 to +1.5. A Z-score equal to 0 means that the score for particular gene is the same as the mean for all conditions, while a Z-score of +1 indicates that the corresponding value is one standard deviation above the mean. Therefore, the variability in histone marks could be cross-compared between genes. Data represent one experiment with two independent biological replicates pooled from 7-9 independent mice.

(C-E) Naïve WT and *Tbx21*$^{+/-}$ LCMV-specific CD4$^+$ Thy1.1$^+$ cells were adoptively transferred into C57BL/6 mice. Recipient mice were infected with LCMV. 10 days after infection, CD4$^+$ Thy1.1$^+$ donor T cells were isolated and re-activated for 2 rounds with weekly re-activation under neutral or Th2 conditions. Subsequently, *in vitro*-re-activated CD4$^+$ Thy1.1$^+$ cells were re-transferred into naïve C57BL/6 mice.

(C) Seventeen days after transfer, total splenocytes from CD4$^+$ Thy1.1$^+$ recipient mice were re-stimulated for 24 h with $GP_{61-80}$ peptide. Cytokine production in the supernatants was measured using CBA.

(D) Seventeen days after transfer, GATA-3 relative expression in $CD4^+$ $Thy1.1^+$ donor cells isolated from spleen was determined. Depicted is GATA-3 relative protein expression in reprogrammed $CD4^+$ $Thy1.1^+$ compared with GATA-3 values in the respective $CD4^+$ $Thy1.1^+$ cells treated under neutral conditions.

(E) GATA-3 expression in $CD4^+$ $Thy1.1^+$ donor cells isolated from spleen 17 days after transfer and cultured for 5 days with $GP_{61\text{-}80}$ and APCs. Geometric mean indices represent the factor of change of the geometric means of the GATA-3 staining compared with the geometric mean values of the respective isotype control staining.

(C, E) Data represent one experiment with n = 4-6 mice per group and time point.

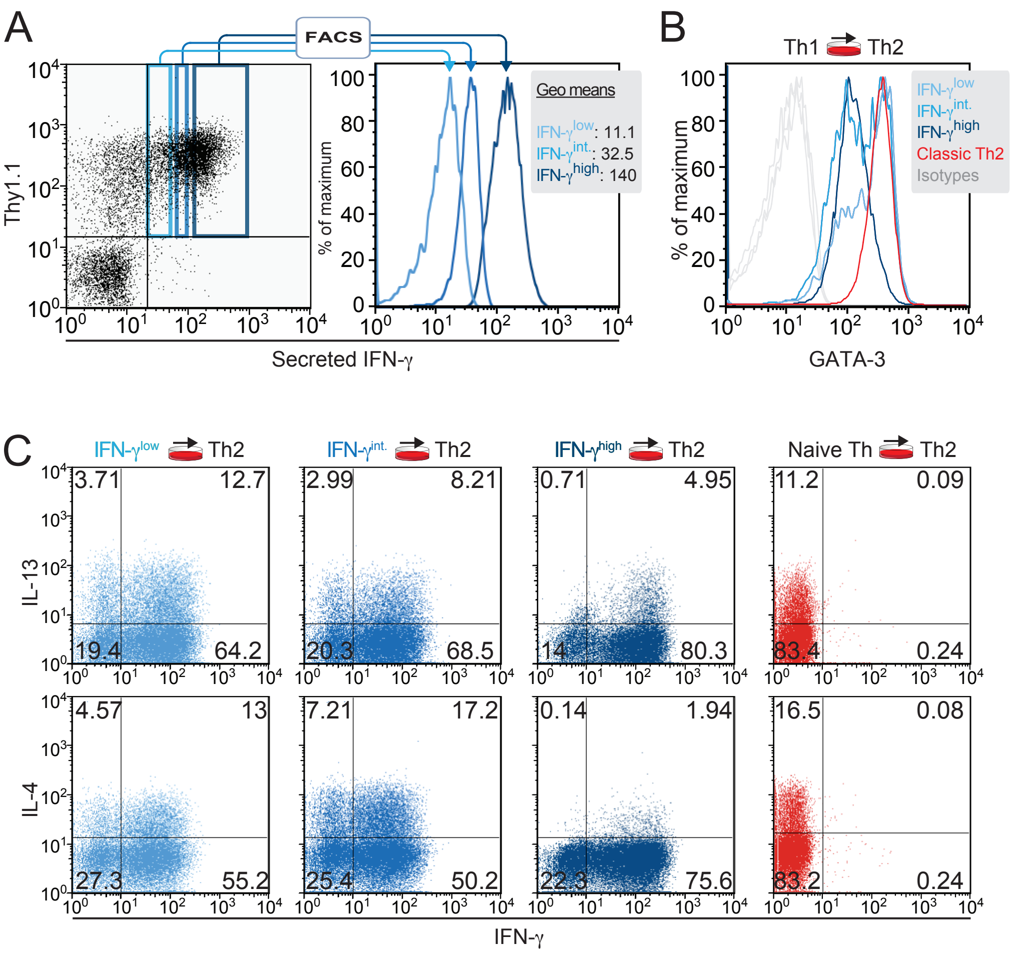
A
FACS
Thy1.1
Secreted IFN-γ
% of maximum
Geo means
IFN-γlow: 11.1
IFN-γint.: 32.5
IFN-γhigh: 140
B
Th1 Th2
IFN-γlow
IFN-γint.
IFN-γhigh
Classic Th2
Isotypes
GATA-3
C
IFN-γlow Th2
IFN-γint. Th2
IFN-γhigh Th2
Naive Th Th2
IL-13
IL-4
IFN-γ
3.71
12.7
19.4
64.2
2.99
8.21
20.3
68.5
0.71
4.95
14
80.3
11.2
0.09
83.4
0.24
4.57
13
27.3
55.2
7.21
17.2
25.4
50.2
0.14
1.94
22.3
75.6
16.5
0.08
83.2
0.24

Figure S6

**Figure S6. Sorting of IFN-γ–secreting effector cells according to their individual IFN-γ expression levels predicts effector Th1 cell plasticity**

(A-C) Naïve LCMV-specific $CD4^+$ $Thy1.1^+$ cells were adoptively transferred into C57BL/6 mice and subsequently infected with LCMV. ns, not significant; * $P < 0.05$; ** $P < 0.01$; *** $P < 0.001$.

(A) On day 10 after infection, LCMV-specific IFN-γ secreting $CD4^+$ $Thy1.1^+$ donor T cells were identified by IFN-γ secretion assay after re-stimulation with $GP_{61\text{-}80}$ peptide and sorted according to their IFN-γ expression levels by FACS. Data represent one experiment, LCMV-specific IFN-γ secreting $CD4^+$ $Thy1.1^+$ donor T cells were sorted from 30 mice.

(B, C) Isolated populations were rested for 5 days in the presence of IL-2, anti–IL-4, anti–IFN-γ, and anti–IL-12. The isolated populations were then reactivated for 1-2 rounds under Th2 conditions. Naïve LCMV-specific $CD4^+$ $Thy1.1^+$ cells reactivated under Th2 conditions served as a control.

(B) GATA-3 protein expression of $CD4^+$ $Thy1.1^+$ cells was analyzed by FACS.

(C) On day 5 of secondary stimulation, cells were reactivated with PMA/ionomycin and stained for CD4 and intracellular cytokines. Percentage of $cytokine^+$ $CD4^+$ T cells is indicated. (B-C) Data represent two independent biological replicated.

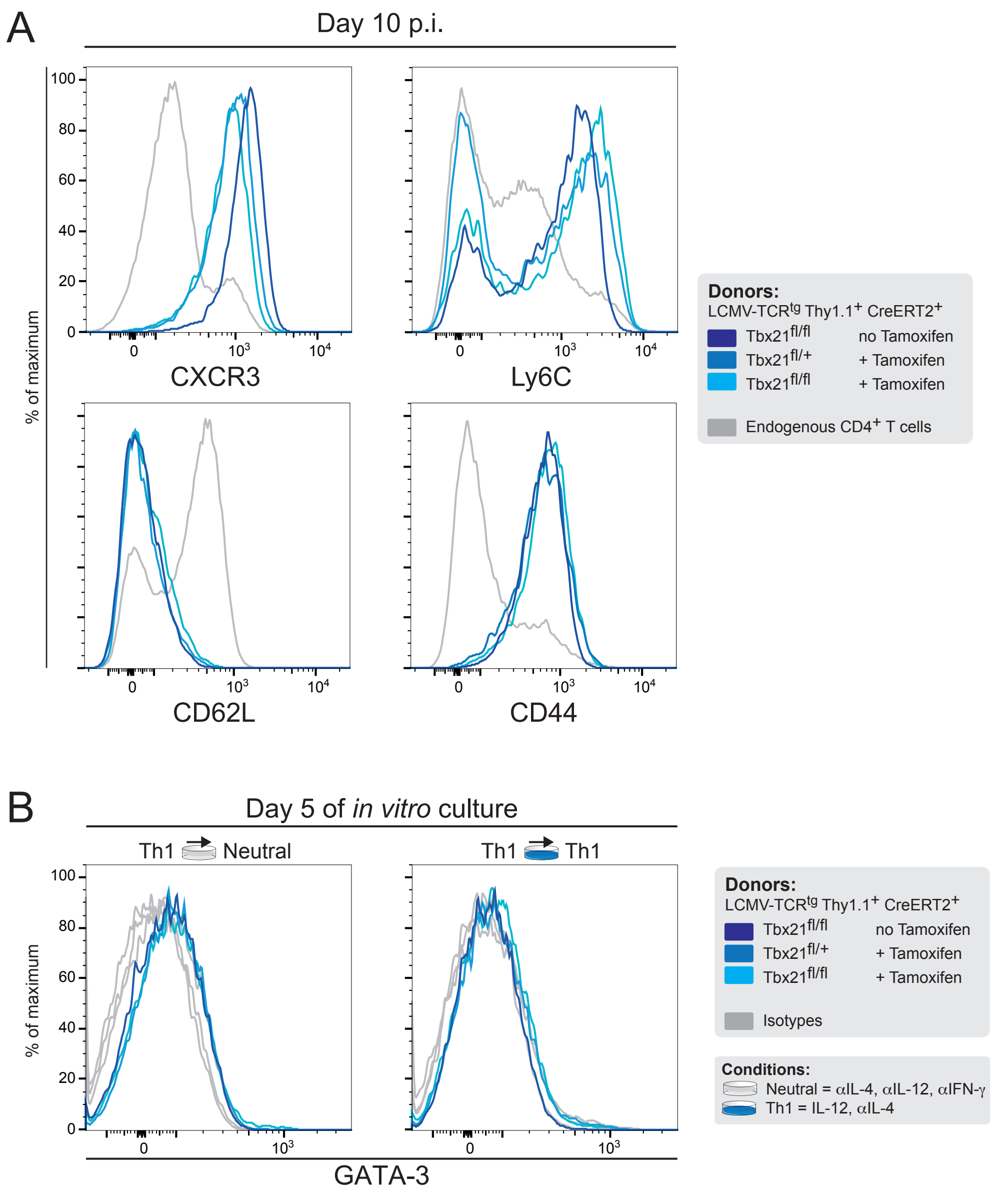

A
Day 10 p.i.
% of maximum
CXCR3
Ly6C
CD62L
CD44
Donors:
LCMV-TCRtg Thy1.1+ CreERT2+
Tbx21fl/fl no Tamoxifen
Tbx21fl/+ + Tamoxifen
Tbx21fl/fl + Tamoxifen
Endogenous CD4+ T cells
B
Day 5 of in vitro culture
Th1 Neutral
Th1 Th1
GATA-3
Isotypes
Conditions:
Neutral = αIL-4, αIL-12, αIFN-γ
Th1 = IL-12, αIL-4


Figure S7

**Figure S7. Continuous T-bet expression safe-guards Th1 cell stability**

Naïve WT, *Tbx21*$^{fl/+}$ and *Tbx21*$^{fl/fl}$ LCMV-specific CD4$^+$ Thy1.1$^+$ cells were adoptively transferred into Th1.2$^+$ *Cre*$^+$ C57BL/6 mice. Recipient mice were infected with LCMV. On day 7, 8, and 9 recipient mice were treated with tamoxifen, and CD4$^+$ Thy1.1$^+$ donor T cells were isolated and re-activated for 2 weeks with re-stimulation after 5 days under neutral (anti–IL-12, anti–IL-4, and anti–IFN-$\gamma$) or Th2 conditions. Th2 cells derived from WT naïve LCMV-specific CD4$^+$ Thy1.1$^+$ cells served as control. ns, not significant; * $P < 0.05$; ** $P < 0.01$; *** $P < 0.001$. Data are representative of two independent experiments.

(A) Surface expression of the indicated proteins was determined on WT, *Tbx21*$^{fl/+}$ and *Tbx21*$^{fl/fl}$ LCMV-specific CD4$^+$ Thy1.1$^+$ cells on day 10 after infection.

(B) GATA-3 protein expression of CD4$^+$ Thy1.1$^+$ cells kept under neutral conditions was analyzed by FACS.

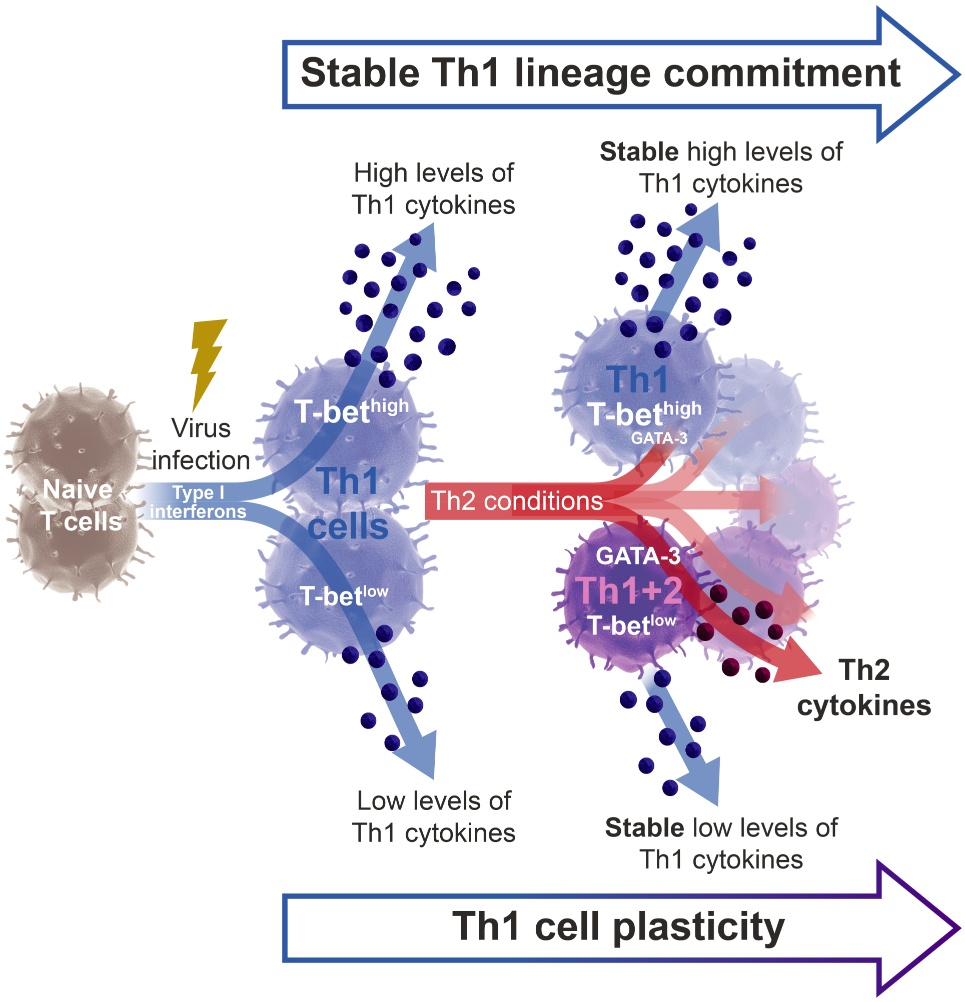
Stable Th1 lineage commitment
High levels of
Th1 cytokines
Stable high levels of
Th1 cytokines
Virus
infection
Naive
T cells
Type I
interferons
T-bet high
Th1
cells
T-bet low
Th1
T-bet high
GATA-3
Th2 conditions
GATA-3
Th1+2
T-bet low
Th2
cytokines
Low levels of
Th1 cytokines
Stable low levels of
Th1 cytokines
Th1 cell plasticity

Figure S8

**Figure S8. T-bet quantities regulate Th1 cell plasticity and commitment**

Graphical abstract summarising the results presented in this study. Type-I interferons triggered by infection modulate T-bet expression states in Th1 cells leading to the generation of a Th1 effector cell pool containing Th1 cells expressing distinct levels of T-bet and IFN-$\gamma$. When subjected to Th2 skewing conditions, these distinct Th1 cells acquire various levels of GATA3 expression depending on their initial T-bet expression levels. The reprogrammed Th1+2 phenotypes exhibit a stable phenotype and a mixed Th1/Th2 epigenetic landscape.

### SUPPLEMENTAL EXPERIMENTAL PROCEDURES

#### Mice

C57BL/6 and C57BL/6 mice congenic for Thy1.1 (B6.PL-Thy1a/CyJ) mice were bred at the Institute of Experimental Medicine (FEM), Charité - University Medicine Berlin, and were used as recipients for adoptive cell transfers at the age of 8-12 wk. *Il12b*$^{-/-}$ mice (Magram et al., 1996), *Ifnar*$^{-/-}$ and *Ifngr1*$^{-/-}$ mice (Muller et al., 1994), *Tbx21*$^{fl/lf}$ mice (Intlekofer et al., 2008), R26-CreERT2 (Ventura et al., 2007) and *Tbx21*$^{-/-}$ mice (Szabo et al., 2002) were purchased from Jackson laboratories (all on a C57BL/6 background). *Il4ra*$^{-/-}$ (Kopf et al., 1993)and Tbet-ZsGreen (Zhu et al., 2012) reporter mice were kindly provided by Manfred Kopf (Institute of Molecular Health Science, Swiss Federal Institute of Technology, Switzerland) and Jinfang Zhu (Cytokine Biology Unit,National Institutes of Health; Zhu et al. 2012), respectively. SMARTA1 TCR-transgenic mice, which express a TCR specific for the LCMV epitope GP61–80 (Oxenius et al., 1998) were crossed to Thy1.1$^{+}$ B6.PL mice to generate TCR$^{tg}$ Thy1.1$^{+}$ mice and then crossed to *Ifngr1*$^{-/-}$ mice, *Ifnar*$^{-/-}$ mice, *Tbx21*$^{-/-}$ mice, T-bet-ZsGreen mice, R26-CreERT2 and *Tbx21*$^{fl/lf}$ mice.

#### Adoptive T cell transfer

Naive LCMV-specific CD4$^{+}$ T cells (1–3 x $10^5$ cells) were given intravenously in 300 µl BSS to yield a seeding of approx. 1–3 x $10^4$ cells per mouse (Hataye et al., 2006). In some experiments where the recipients were left uninfected, 0.5-1 x $10^7$ cells were transferred to yield a seeding of approx. 0.5-1 x $10^6$ cells per mouse.

#### Viruses

LCMV-ARM and LCMV-WE strains were propagated on BHK21 or L929 cells respectively. Mice were infected intravenously with 200 PFU.

**T cell activation and differentiation**

Naive MACS sorted LCMV-TCRtg $CD4^+$ T cells ($CD4^+$ $CD62L^+$ $CD25^-$) from spleens and lymph nodes isolated from the above-mentioned mice were cultured in RPMI 1640 (IMEM for Th17 conditions) supplemented with 10% (vol/vol) FCS (Gibco), L-glutamine (2µM; Gibco), penicillin (100U/ml; Gibco), streptomycin (100µg/ml; Gibco) and β-mercaptoethanol (50nM; Sigma) in the presence of APCs and 1 µg/ml GP64–80 peptide (Neosystem) together with different combinations of antibodies and cytokines: for Th1 differentiation, 3 ng/ml IL-12 (R&D Systems), 5 ng/ml IL-2 (Peprotech), and 10 µg/ml anti–IL-4 (11B11); for Th2 differentiation, 30 ng/ml IL-4 (Sigma-Aldrich), 5 ng/ml IL-2 (Peprotech), 10 µg/ml anti–IL-12 (C17.8), and 10 µg/ml anti–IFN-γ (AN18.17.24); for neutral conditions, 5 ng/ml IL-2 (Peprotech), 10 µg/ml anti–IL-4 (11B11), 10 µg/ml anti–IL-12 (C17.8), and 10 µg/ml anti–IFN-γ (AN18.17.24) (Hegazy et al., 2010); for Th9 differentiation, 20 ng/ml IL-4 (Sigma-Aldrich), 3 ng/ml TGFβ (Peprotech), 5 ng/ml IL-2 (Peprotech), 10 µg/ml anti–IL-12 (C17.8) and 10 µg/ml anti–IFN-γ (AN18.17.24); for Treg differentiation, 5 ng/ml TGFβ (Peprotech), 5 ng/ml IL-2 (Peprotech), 10 µg/ml anti–IL-4 (11B11), 10 µg/ml anti–IL-12 (C17.8) and 10 µg/ml anti–IFN-γ (AN18.17.24); for Tfh conditions, 20 ng/ml IL-6, 20 ng/ml IL-21, 10 µg/ml anti–IL-4 (11B11), 10 µg/ml anti–IL-12 (C17.8) and 10 µg/ml anti–IFN-γ (AN18.17.24); for Th17 conditions, 20 ng/ml IL-6 (Peprotech), 20 ng/ml IL-23 (Peprotech), 1 ng/ml TGFβ (Peprotech), 10 µg/ml anti–IL-12 (C17.8), and 10 µg/ml anti–IFN-γ (AN18.17.24). Cell cultures were split on days 2 and 4. On day 6, cells were reactivated with fresh Thy1.2-depleted C57BL/6 splenocytes plus GP64–80 peptide, 5 ng/ml IL-2 (R&D Systems), cytokines and anti-cytokine mAbs as above.

**Sorting of live IFN-γ–secreting cells**

The cytometric cytokine secretion assay was performed as described before (Assenmacher et al., 1998; Löhning, 2003). Briefly, cells were restimulated for 2.5 h to 4 h, followed by labeling with an IFN-γ–specific capture matrix (Miltenyi Biotec). Control samples were either incubated with the respective recombinant cytokine to control the homogeneous labeling of all cells with the cytokine capture matrix (high control), or they were put immediately in ice-cold buffer to prevent cytokine secretion (low control). The capture matrix-labelled cells were kept in 37°C warm medium for 20 min. Matrix-captured cytokine was stained on the cell surface with anti–IFN-γ-PE (Miltenyi Biotec). Cells were subsequently FACS sorted according to their PE signal intensity.

**Flow cytometry and cell sorting**

Lymphocytes were isolated from peripheral blood, spleen and lymph nodes as described previously(Hegazy et al., 2010). Peripheral blood lymphocytes (PBLs) were purified by underlaying Histopque 1083 (Sigma-Aldrich) and performing density centrifugation (400g at 20°C for 30 min). Adoptively transferred T cells were detected and analyzed with mAbs to Thy1.1 (OX-7), Thy1.2 (53-2.1), CD4 (RM4-5), CD62L (MEL-14), CD44 (IM7), IL-7Rα (A7R34), CCR4 (2G12), CXCR3 (CXCR3-173), PD-1 (J43), CCR6 (29-2L17), CXCR5 (J252D4), IL33R (DJ8), IL18R (BG/IL18RA). To prevent unspecific binding of mAb, all samples were preincubated with 10 µg/ml of blocking anti-FcεRII/III (2.4G2; ATCC) and 50 µg/ml of purified rat IgG (Biotrend). For FACS sorting experiments, samples were sorted on a FACSAria II. Samples were acquired on FACSCaliburTM, FACSCanto II or FACS LSRFortessa (Becton Dickinson), and data were analyzed with FlowJo (Tree Star). In some experiments, samples were enriched for $CD4^+$ T cells. Spleen and lymph node cells were stained ex vivo with FITC-conjugated anti-CD4, and subsequently incubated with anti-FITC microbeads (Miltenyi). $CD4^+$ T cells were enriched on LS columns (Miltenyi).

**Intracellular cytokine staining**

For analysis of intracellular cytokines, spleen or LN cells were restimulated with either GP64-80 (1 µg/ml) (Neosystem) or with PMA (5 ng/ml) and ionomycin (500 ng/ml; Sigma). 5 µg/ml brefeldin A (Sigma-Aldrich) was added at 2 h. After 4–5 h, cells were fixed with 2% formaldehyde (Merck) and stained with anti-CD4 or anti-CD8 and anti-Thy1.1 or anti-Thy1.2, and with the following rat anti–mouse cytokine mAbs or isotype control mAbs in permeabilization buffer containing 0.05% saponin (Sigma-Aldrich) as described previously (Hegazy et al., 2010): anti–IFN-γ (XMG1.2), anti–TNF-α (MP6-XT22), anti–IL-2 (JES6-5H4), anti–IL-4 (11B11), anti–IL-5 (TRFK5), anti–IL-9 (RM9A4), anti–IL-10 (JES5-16E3), anti–IL-13 (38213.11), anti–IL-17A (TC11-18H10.1), anti–IL-22 (1H8PWSR), and anti–GM-CSF (MP1-22E9). Rat IgG1 (R3-34) and rat IgG2a (R35-95) isotype control mAbs were used at the same concentrations as the respective anti-cytokine mAbs.

**Transcription factors staining**

T-bet, GATA-3, FoxP3, RORγt, and BCL-6 protein expression was analyzed using the FoxP3 staining buffer set (eBioscience) according to the manufacturer's instructions. Briefly, prior to fixation, cells were stained with PerCP-conjugated anti-CD4 and Pacific Blue-conjugated anti-Thy1.1. Cells were then fixed with 1x Fixation/Permeabilization buffer, followed by intracellular staining with fluorochrome-labeled antibodies against the above-mentioned transcription factors in 1x permeabilization buffer. Cells were washed with 1x permeabilization buffer and immediately analyzed. PE-conjugated anti-Tbet (4B10), Alexa-647-conjugated anti-GATA-3 (TWJA) and FITC-conjugated anti-FoxP3 (150D/E4) were purchased from eBioscience. PE-conjugated anti-BCL6 (K112-91) and RORγt (Q31-378) were purchased from BDBioscience.

**Intracellular STAT protein staining**

Intracellular STAT6 stainings were done using Phosphflow buffers according to manufacturer's instructions (BD Biosciences). Briefly, $CD4^+$ T cells were stimulated with the indicated cytokines or left untreated at 1-2 x $10^7$ cells/ml in serum free RPMI at 37 °C after serum starvation period of 3 h, stimulation was stopped with ice-cold PBS. Cells were fixed with pre-warmed 1x BD™ Phosflow Lyse/Fix Buffer (BD Biosciences) for 10 min at 37 °C. The cells were permeabilized with ice cold BD™ Phosflow Perm Buffer III (BD Biosciences) for 30 min on ice. Subsequently, the cells were stained in PBS with 0,2 % (vol/vol) BSA for 30 min with PerCP-conjugated anti-CD4, Pacific Blue-conjugated anti-Thy1.1 and PE-conjugated anti-pSTAT6 (38/p-Stat4), Alexa-647-conjugated anti-STAT6 (23/Stat6) (all from BD Biosciences). Cells were washed and immediately analyzed. pSTAT6 data are represented as fold increase (factor of change compared with untreated control). Total STAT6 data are represented as Geo Mean index (factor of change compared with isotype control staining).

**Cytokine analysis in supernatants**

The concentration of IL-4, IL-5, IL-13, IL-10 and IFN-$\gamma$ in supernatants were determined with Cytometric Bead Arrays (BD Biosciences) according to the manufacturer's instructions.

**RNA isolation and quantitative PCR**

Total RNA was extracted from cells with RNAeasy Mini Kit according to the manufacturer`s instructions (Qiagen). RNA was reverse transcribed to cDNA using random primers and the First Strand cDNA Synthesis Kit (Fermentas). Real-time PCR was performed with Taqman Fast Universal PCR Mastermix (Life Tech) using the Taqman GeneExpression Assays. Each assay was performed in duplicates with a Real-Time PCR Detection System (BIORAD, CFX96) and gene expression levels were normalized to *Gapdh* or *Actin* and then normalized to $CD4^+$ $Cd44^{low}$ $CD62L^+$ naïve T cells.

**Chromatin Immunoprecipitation and Sequencing (ChIP-Seq)**

WT, *Tbx21*$^{+/-}$, *Tbx21*$^{-/-}$ LCMV-specific $CD4^+$ $Thy1.1^+$ cells from LCMV challenged mice were isolated and MACS sorted on day 10 of infection. Cells were culture for two rounds under neutral or Th2 conditions as described above. On day 4 of culture the cells were harvested and processed for Chip-Seq. Cells were incubated for 10min at room temperature with 1% formaldehyde. Cross-linking was terminated by addition of glycine to a final concentration of 125mM. ChIP was performed using iDeal ChIP-seq Kit (Diagenode) according to the manufacturer's instructions and as previously described. Antibodies against H3K4me3 (ab8580, Abcam), H3K4me1 (ab8895, Abcam), H3K27ac (ab4729, Abcam), H3K27me3 (ab6002, Abcam) and H3K9me3 (ab8898, Abcam) were used.

Library preparation for ChIP-seq was performed using NEBNext ChIP-Seq library Prep Master Mix set kit. Two biological replicates per condition were sequenced using Illumina HiSeq 2000 (50bp single-end) sequencing technology. 12 indexes were used per sequencing lane.

**Library preparation for Illumina**

10ng of chromatin-immunoprecipitated DNA was end-repaired and dA-tailed. Subsequently, adaptor was ligated to the dA-Tailed DNA. DNA was cleaned up using AMpure XP beads after each step. Adaptor-ligated DNA was selected size with AMpure XP beads using different DNA to beads ratio to an average fragment size of 200bp. At last step, size-selected and adaptor-ligated DNA was enriched by PCR with different index primers and cleaned up using AMpure XP beads. Purified library DNA was quantified with Qubit 2.0 and fragment size was assessed using Bioanalyzer (Agilent high sensitivity chip). The raw data from our RNAseq data are deposited at

ArrayExpress (http://www.ebi.ac.uk/arrayexpress/) with accession number E-MTAB-2351.

**ChipSeq data analysis and bioinformatical analysis**

The Chip DNA fragments where sequenced on Illumina HiSeq 2000 sequencing platform (Illumina USA, San Diego). Each of two biological replicates was sequenced twice. To reduce the risk of typing and copy-paste errors the data analysis pipeline was tighten together using Perl (http://www.perl.org/) and optimized to run on a computational cluster.

The quality control of raw data performed with a FastQC- a quality control tool for high throughput sequence data (http://www.bioinformatics.babraham.ac.uk/projects/fastqc). Then reads where mapped using publicly available software Bowtie (http://bowtie-bio.sourceforge.net/index.shtml). For follow-up analysis resulting BAM files are transformed first into BAM format and then into BED format using samtools (http://samtools.sourceforge.net/). Only unique reads where selected for a follow-up analysis. After additional quality control of a uniquely mapped data, both technical and biological replicates were merged together. Based on merged BED files new IGV tracks were calculated with IGVtools (http://www.broadinstitute.org/igv/igvtools) and visualized with Integrative Genomics Viewer (IGV).

To quantify changes in the epigenetic status of T cells related to a T-bet expression we selected a group of genes involved in T cell regulation or immune response, such as cytokines, T cell receptors or transcription factors (full list see in supplement materials Table 1). For each selected genes and each sample a normalized mean occupancy was calculated: total number of sequencing reads in the transcript and in 2 adjacent flanking regions of 5kb each was calculated and divided by transcript length and total number of reads in merged sample. This value is used to represent the epigenetic status of the corresponding gene under different experimental conditions.

**RNA Sequencing library preparation**

RNA was extracted from $10^6$ cells using RNeasy mini kit (Qiagen) with a separate step for DNAse I digestion (Qiagen). ERCC Spike-in control mix-1 (Life technologies) was added to each sample prior to processing (1µL of a 1:10 dilution of mix-1). The rRNA depletion and library construction was processed with the ScriptSeq Complete Gold Kit (Epicentre) according to the manufacturers' instructions. The index for individual samples was added in order to multiplex 9 samples in each sequencing lane. The quality of libraries was assessed using a DNA1000 chip on the Bioanalyzer (Agilent) and the concentration was measured using the Qubit DNA assay. Sequencing was performed on the Illumina platform with HiSeq 2000 paired- end 100 bp sequence type.

**Postprocessing of RNA-Seq data**

Sequencing reads were aligned to the mouse genome (GRCm38.p4 build) using STAR (v2.4.0j). Read counts for each transcript were determined as the total number of reads mapped to unambiguous exons using htseq-count (v0.6.1). For determining differentially upregulated genes between Th1 and Th2 conditions we used the function DESeq() from DESeq2 (v1.20). The genes were selected according to the multiple-testing-corrected p-value (according to Benjamini-Hochberg) and the largest logarithmic fold-change values (logFC).

**Statistical analysis**

Two groups were compared using a two-tailed unpaired Student's *t*-test. More than two groups were compared using one-way ANOVA with Bonferroni's post-test for multiple comparisons. Time courses of multiple groups were compared with two-way ANOVA.
